# A Time-Resolved Single-Cell Atlas Reveals Infection-Status, Age-, and Sex-Dependent Immune Responses Drive Viral Disease Severity

**DOI:** 10.64898/2026.08.10.744019

**Authors:** Lujia Chen, Aodong Qiu, Jooyoung Kim, Nebal AbuHusein, Mengyao Lu, Karen Agaronyan, Kexin Sun, Yifan Yuan, Amy Zhao, Gagan Heda, Xinghua Lu, Georgios Kitsios, Alicia N Rizzo, William Bain, Toru Nyunoya, Tomeka Suber, John Evankovich, Faraaz Shah, Charels Dela Cruz, Edward Manning, Lokesh Sharma

**Author notes:** Correspondence: Lokesh Sharma. contributed equally (Lujia Chen, Aodong Qiu, Jooyoung Kim).

## Abstract

The majority of mortality during viral infections occurs in older males; however, underlying mechanisms by which age and sex shape antiviral immunity and pathological inflammatory responses remain incompletely understood. Here, we performed time-resolved single-cell RNA sequencing across 16 conditions spanning age, sex, and four stages of influenza infection in mice, generating a high-resolution atlas. Aged mice demonstrate delayed antiviral and inflammatory responses in multiple myeloid cells, impairing viral clearance, which delays recovery. Similarly, endothelial cells from aging mice show prolonged inflammatory and antiviral gene signatures. Altered gene signatures in immune and endothelial cells result in a shift in endothelial-immune interactions in the aged lung. Further, the infection status of the cell is a major driver of transcriptional state, with infected myeloid cells exhibiting broad upregulation of genes, including interferon-stimulated, inflammatory, complement, and oxidative stress-related genes. To assess whether these age-associated transcriptional patterns are conserved in humans, we examined BAL cells obtained from healthy individuals and COVID-19 patients, and found that immune cells from aged COVID-19 patients had elevated antiviral and pro-inflammatory gene expression compared to cells from young patients. Our analyses of sex differences identified that multiple myeloid cell types in aged male mice, but not in young male mice, show persistent inflammatory responses at later stages of infection, a likely mechanism contributing to elevated mortality in older males. These data reveal how infection status of the cell, age, and sex interact to drive persistent inflammation and impaired resolution, providing a foundational resource for designing age- and sex-specific therapeutic strategies.

## Introduction

Aging represents the most significant risk factor for viral infection-mediated mortality in humans. Over 95% of the mortality related to both seasonal and pandemic influenza occurs in those over 65 years [1, 2]. Young females show high susceptibility to influenza infection and elevated adverse effects to influenza vaccines [3, 4]; however, the majority of mortality occurs in older males in influenza and other respiratory viral diseases such as COVID-19 [5, 6]. The age-dependent increase in susceptibility to influenza infection is multifactorial, including increased comorbidities, impaired innate immune responses (e.g., by macrophages and dendritic cells), and decreased adaptive immune responses measured by antibody titers [7, 8]. Further, aged lungs have impaired ability to repair the tissue damage caused by the viral infections, contributing to the prolonged illness and impaired recovery.

Enhanced susceptibility to influenza infection in aged hosts can be validated in mouse models where multiple components of aging contribute to immune impairment against influenza infections [9–11]. Prior research has also identified female sex-associated susceptibility to influenza infections using mouse models [12]. However, a comprehensive understanding of age and sex-specific mechanisms of influenza susceptibility at the single-cell level, during different stages of influenza infections, remains unclear.

First, we confirmed age-dependent susceptibility to influenza infection in mice. We then performed a comprehensive single-cell RNA sequencing of mouse lungs infected with influenza virus at baseline (day 0), at the initial stage of infection (day 3), at the peak inflammatory/injury stage (day 7), and at the repair phase (day 11) to compare the disease kinetics between young and aged male and female mice at the single-cell level. Our data provide the first comprehensive single-cell RNA sequencing analysis to determine host response to influenza infection in an age-, sex-, and disease stage-dependent manner. Our findings reveal that persistent viral presence is the key driver of inflammatory responses in the myeloid cell compartment, a likely mechanism explaining exacerbated inflammatory injury and impairing tissue repair in aged mice. These findings are supported by publicly available human BAL single-cell RNA sequencing data from COVID-19 patients.

## Materials and Methods

### Mouse Models

All animal experiments were approved by Yale IACUC. Young (8-12 weeks) and aged (>20 months) C57BL/6 mice were used for the experiments. Influenza infections were performed using H1N1/PR8 strain with 5 PFUs/mouse suspended in 50 μL PBS under ketamine/xylazine anesthesia by intranasal route as described previously [13, 14].

### Flow Cytometry

White blood cell counts (WBC) were counted using a Coulter counter as published previously [14, 15]. Flow cytometry was performed with multicolor flow cytometry protocols similar to those published previously [16].

### Histology

Lung histology was prepared by inflating the lungs with 0.5% low-melting agar at 22 cm height of column. Lungs were then fixed in 4% paraformaldehyde. Lungs were then embedded, sectioned, and stained with hematoxylin and eosin.

### Single-Cell Suspension and 10X Sequencing

Lung samples obtained from young and aged mice at different stages of influenza infection were harvested after flushing the lung with sterile PBS by cardiac puncture. Lungs were chopped into small pieces and were digested with collagenase type 4 (2 mg/ml) and DNase (20 U/mL) in DMEM with 10 % FBS to prepare a single-cell suspension. We performed magnetic-activated cell-sorting using CD45+ beads. A 50:50 mixture of CD45+ and CD45- cells was prepared to enrich for non-immune cells. We used the 10X Genomics Chromium platform (3’ v3.1 kit), a droplet-based microfluidic system, to barcode unique mRNA molecules of each cell. We then performed reverse transcription, cDNA amplification, fragmentation, adaptor ligation, and sample index PCR according to the manufacturer’s protocol. We evaluated for quality control by tracing cDNA after barcoding using a high-sensitivity bioanalyzer. Our core facility sequenced the final cDNA libraries on a HiSeq 4000 Illumina platform, aiming for 150 million paired-end reads per library, at the manufacturer’s recommended read 1 and read 2 lengths. Raw sequencing reads were demultiplexed based on sample index adaptors, which were added during the last step of cDNA library preparation.

### Data Processing

The raw single-cell RNA sequencing data were preprocessed using CellRanger (v7.2.0), including aligning reads to the reference genome, and generating cell by gene expression count matrices. To identify viral reads in the sequencing data, the influenza A genomic reference was created by combining the mouse reference library with a H1N1 infection genome reference, including the following genes: PB2, PB1, PB1-F2, PA, PA-X, HA, NP, M2, M1, NS2, and NS1. Count matrix for each sample was generated by running CellRanger count function on the FASTQ files using default parameters and customized reference genome. Cells expressing at least one infection gene were labeled as “infected”, which was added to the single-cell object’s metadata.

The data filtration, normalization, scaling, integration, and clustering were performed using Seurat (v5.1.0) in the R environment. The cells were filtered to remove mitochondrial genes, debris and doublets. Cells passing quality control (QC) from each sample were integrated for analysis using Seurat functions FindIntegrationAnchors and IntegrateData [17, 18]. Ambient RNA contamination was removed using DecontX (v1.20.0)[19] from the celda R package. Raw count matrices were extracted from the RNA assay and passed to the decontX function along with pre-computed cluster assignments to guide the contamination estimation. Cells with a contamination fraction exceeding 0.5 were excluded. The counts were normalized to the total counts and multiplied by a 10,000 scaling factor. Principal component analysis (PCA) was performed on the 2,000 identified highly variable genes for dimensionality reduction, retaining 20 principal components prior to cell clustering. Batch effects across samples were corrected using Harmony (v0.1.1) [20]. Cell clustering was performed on the integrated data using a shared nearest-neighbor (SNN) graph–based approach implemented with the FindNeighbors function in Seurat, followed by modularity optimization using the Louvain algorithm with the FindClusters function (resolution = 1.5). All cells were subsequently embedded into two-dimensional Uniform Manifold Approximation and Projection (UMAP) space. Top marker genes for each identified cell cluster were identified using the FindMarkers function and manually matched to canonical cell types for annotation.

### Data Analysis

To perform the differential expression analysis, we utilized the Wilcoxon rank-sum test with a logfc.threshold of 0.4 and a min.pct of 0.01, employing the FindMarkers function from Seurat. The AddModuleScore function from Seurat was applied to calculate the functional gene set of interest. The influenza signature includes 30 genes as described in our prior publication [14], and the hallmark inflammatory signature was obtained from the GSEA hallmark cohort (GSEA M5932). Wilcoxon test was used to compare module scores between the conditions of interest, and p-values were adjusted using the Benjamini-Hochberg procedure to correct for false discovery rate (FDR). Enriched pathway analysis was performed using the R package “fgsea” v1.32.4 [21]. Trajectory analysis was conducted using Monocle3 (v1.3.7) [22].

### Human COVID-19 Cohort

Publicly available human single-cell RNA sequencing data from the BAL samples were obtained from previously published studies [23, 24]. The patient population was divided in young (<40 years) and aged (>65 years) groups to compare the gene signature. Baseline young and old samples were obtained from the study [25]. The COVID-19 Response Score (CRS) was calculated as described previously [26]. This dataset provided us with BAL samples from 39 young (225439 cells) and 17 aged healthy (63680 cells) individuals, along with 9 samples from 4 young COVID-19 patients (65554 cells) and 9 samples from 6 aged COVID-19 patients (45122 cells).

## Results

### Influenza infection causes severe disease and impairs recovery in the aged host

To validate our mouse model of age-mediated susceptibility to influenza, we infected young (8- 12 weeks) and aged (>20 months) mice of both sexes. Mice were monitored daily for body weight changes for up to 16 days. Young male mice lost minimal body weight (Fig. 1A).

**Fig. 1.**
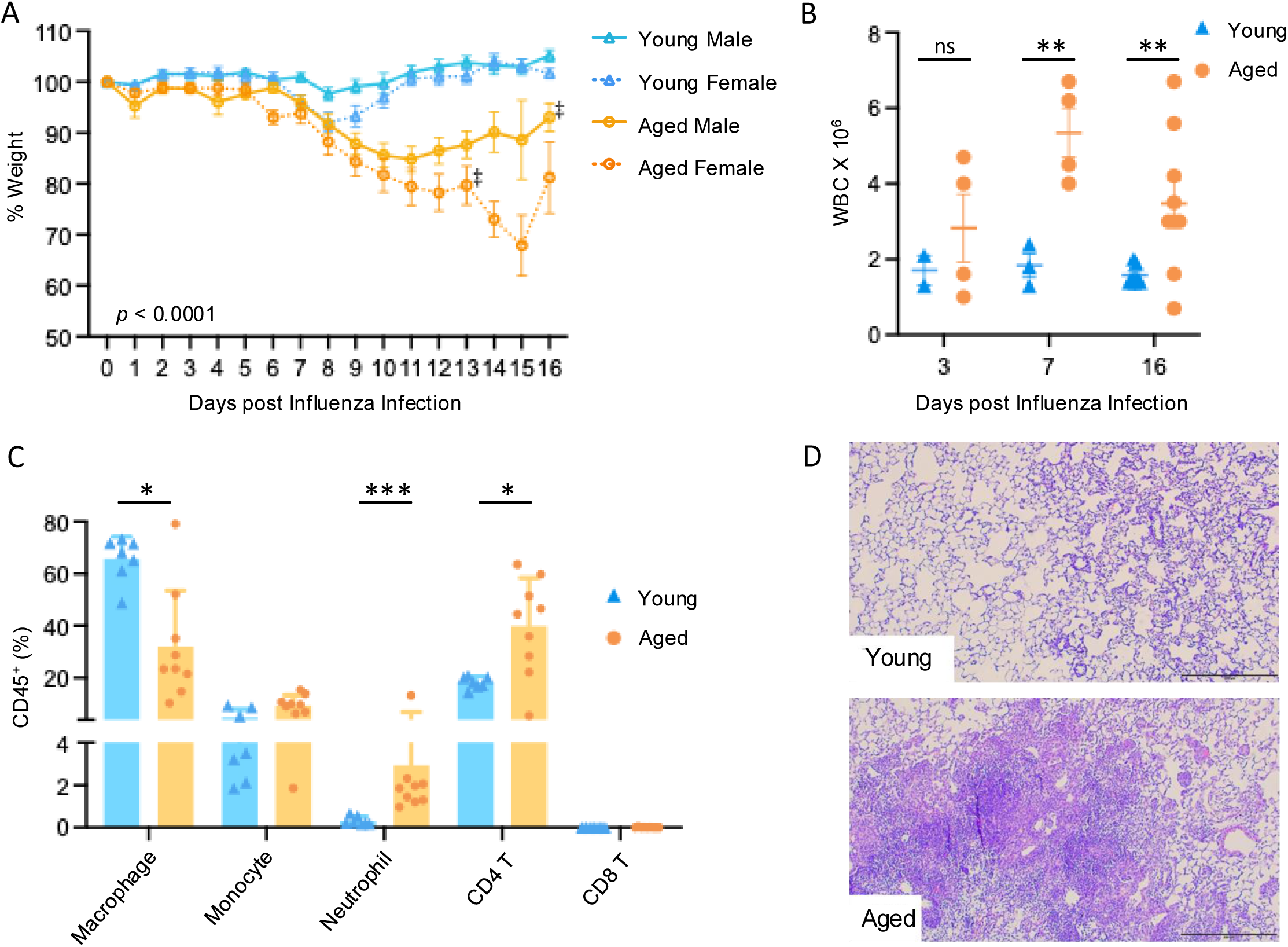
Aging renders mice susceptible to influenza infection. Mice were infected with influenza, and their weight changes were measured every day up to 16 days (A). Mice were euthanized at different time points post-infection with influenza, and recruited cells were counted in the bronchoalveolar lavage (BAL) (B). Flow cytometry analysis was conducted to identify the cell types in the BAL on day 16 post-infection (C). Hematoxylin and eosin staining of the lung tissue sections shown at day 16 post-infection (D). The scale bar shown at the bottom is 200 um. N=9-10/group for young and 10-11/group aged mice, each pooled from 3 independent experiments for A. Each dot represents an individual mouse for B. N= 7-9/group for day 16 data shown in Fig. 1B -D. ‡ indicates the mortality of the mouse on the given day. Data were analyzed using unpaired Student’s t-tests or Wilcoxon rank-sum test based on variance between groups, with Holm-Šídák correction for multiple comparisons. Statistical significance was set at p < 0.05. Data are presented as mean ± standard deviation (SD) with individual data points shown.

Compared to male mice, young female mice lost more weight, especially between day 7-9, when influenza disease severity peaks. However, young mice, regardless of sex, recovered their body weights by day 11. In contrast to the young mice, aged mice lost significantly more weight and failed to recover to baseline body weight during the observation period.

We measured inflammatory cell accumulation in the broncho-alveolar lavage (BAL) fluid. Aged mice had elevated inflammatory cell accumulation measured as total WBC counts in the BAL as early as day 7 post-infection and stayed elevated until day 16 of infection (Fig. 1B). Neutrophils and CD4+ T cells show higher accumulation in the aged mice while a decrease in the levels of macrophages was observed (Fig. 1C). Consistent with elevated inflammatory responses in the BAL, we observed a severe pathology in the lung of aged mice, including increased consolidation (Fig. 1D). Collectively, aged mice have elevated inflammatory response, increased pathology, and impaired recovery post-influenza infection.

### Extensive viral replication across multiple myeloid cell types in the aged lung

To understand the underlying mechanisms that drive inflammatory tissue injury and impaired recovery in the aged host, we performed single-cell RNA sequencing of lung tissues. To determine the effects of age and sex on the specific phases of infection, we collected samples at early phase (day 3), peak inflammatory/injury phase (day 7), and resolution phase (day 11) of influenza infection, along with mock-infected controls, designated as day 0. We pooled two biological samples for each time point for each sex, making n = 4 biological replicates each time point for each age group, except day 11 female, which was only obtained from one female mouse. In total, 134,300 cells were obtained from 31 mice. Cell identity was established using cell-specific gene expression markers (Fig. 2A). A Uniform Manifold Approximation and Projection (UMAP) was generated using gene expression profile (Fig. 2B). We identified the expected cell types from the lung, including epithelium, endothelium, interstitial and alveolar macrophages, monocytes, neutrophils, T, and B cells, among others. The percentage of each cell population during infection is indicated in Sup Fig. 1.

**Fig. 2.**
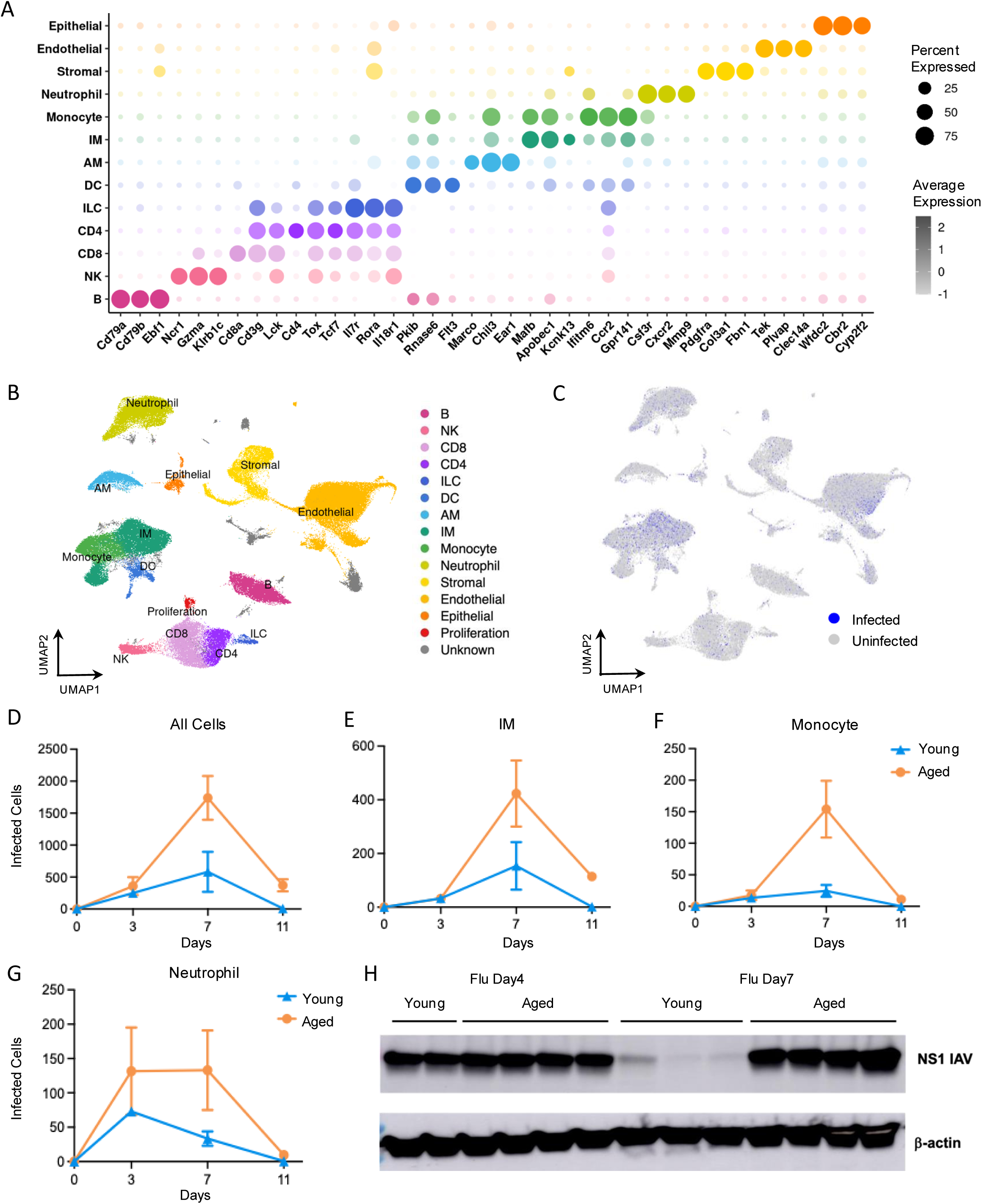
Aging results in extensive viral replication in multiple myeloid cells. Single-cell RNA sequencing was performed on the lung tissue from different time points post-influenza infection. A dot plot demonstrating the gene expression associated with the cell identity (A). A UMAP demonstrating cell identity of the cells obtained from the lung tissue (B). Influenza-positive cells were identified using the expression of influenza genes after DecontX correction (C). Number of total influenza-positive cells (D), interstitial macrophages (E), monocytes (F), and neutrophils (G) between young and aged mouse lungs. Western blot analysis of influenza NS1 protein between young and aged lungs on day 3 and day 7 post-infection (H). β-actin was used as a housekeeping control.

We have previously shown that influenza transcripts render infected cells inflammatory, leading to lung pathology and impaired recovery [14]. To investigate whether lung cells from aged mice have increased susceptibility to infection, we quantified influenza viral transcripts across all pulmonary cell populations. Environmental contamination was removed using Decontx [19].

Our data show that multiple cells including epithelial, endothelial, myeloid, and lymphocytes, contained viral transcripts (Fig. 2C). On day 3, a similar number of infected cells were identified between young and aged mice; however, by day 7, aged mice had a significantly higher number of infected cells compared to the young mice (Fig. 2D). The young mice cleared influenza virus from their lung cells by day 11, while aged mice still had infected cells on day 11 (Fig. 2D).

Myeloid cells such as interstitial macrophages (Fig. 2E), monocytes (Fig. 2F), and neutrophils (Fig. 2G) from aged mice harbored a higher number of viral transcripts compared to young mice. To confirm the elevated viral load in the aged mice, we performed Western blot analysis using independent biological samples collected on day 3 and day 7, confirming higher levels of influenza NS1 protein on day 7 but not on day 3 post-infection in aged mice (Fig. 2H). Taken together, these data show that aging impairs viral clearance, leading to elevated and prolonged presence of viral PAMPs in multiple cell types.

### Neutrophils mount an age and disease stage-specific immune responses during influenza infection

Neutrophil-mediated inflammatory responses are critically important in viral clearance and tissue repair [27, 28]; however, excessive neutrophilic inflammation is associated with lung pathology during influenza infection [29]. To determine aging’s impact on neutrophil responses during influenza infection, we investigated transcriptional changes in neutrophils between young and aged mice that included 14,430 neutrophils across the time course of influenza infection.

Neutrophil gene expression kinetics across infection, shown as top 20 DEGs per time point, are presented in Fig. 3A. Clusters 2, 1 and 4 include genes that peak at day 3, 7 and 11 days post- infection.

**Fig. 3.**
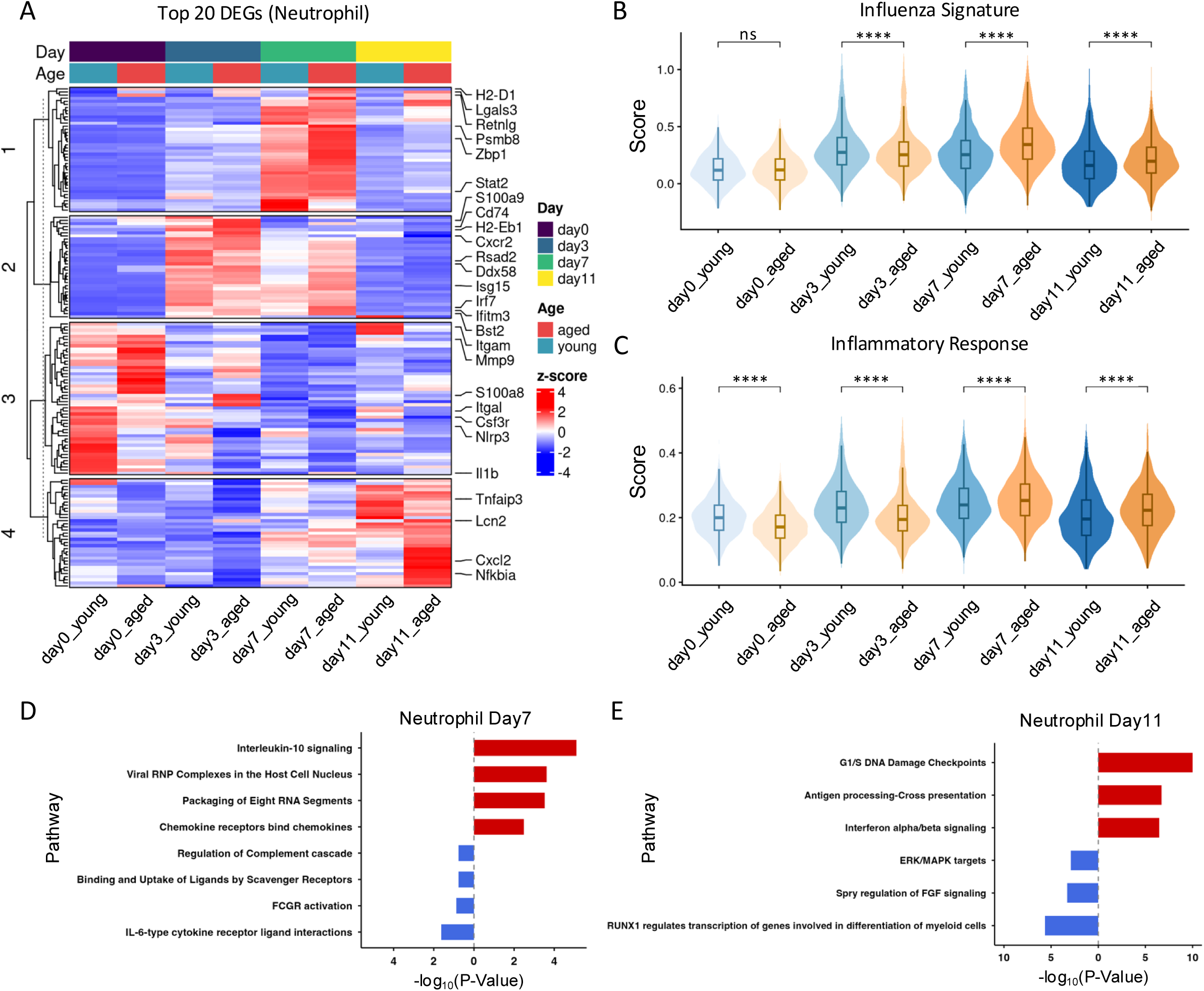
Aging alters neutrophil response to influenza infection in the lung. A heatmap generated using the top 20 unique genes expressed in each condition of age and time post- infection (A). Influenza Signature Score (B) and Inflammatory Response Score (C) during the course of influenza infection between young and aged neutrophils. Pathway analysis was performed on day 7 (D) and day 11 (E) between neutrophils obtained from young and aged mice. IRS and ISS scores were compared using the Wilcoxon rank-sum test. *, P < 0.05; **, P < 0.01; ****, P < 0.0001; ns, not significant.

In mock-infected animals, aged neutrophils had higher basal expression of granulocytic activation genes, including Tnfaip2, Il1b, S100a1/a6/a8/a9/a11, Lyz2, Lcn2, Lgals3, Retnlg, Tyrobp, NADPH-oxidase components, including Cyba and Cybb. On the other hand, young neutrophils had higher expression of genes related to inflammatory response and negative regulators of inflammation including Nfkb1/2, Nfkbid, Nfkbie, Nfkbiz, Tnfaip2/3, Cxcr4, Csf3r, and Ccrl2, suggesting the ability of young cells to exert robust immune response. The interferon-stimulated gene (ISG) signature remained mixed at this time point, where aged neutrophils had elevated Ifitm3, Ifi27l2a, Ifi27, Slfn1 and Slfn2 while the young neutrophils had higher expression of Ifitm3 and interferon receptor Ifnar1.

On day 3 post-infection, young neutrophils had upregulation of chemokine/cytokine response genes, including Il1b, Il15, Nlrp3, Cxcl2, Ccrl2, Pde4b, Ptgs2, Icam1, Tnfrsf1a, Tnfrsf1b and Gpr132. At the same time, young neutrophils had elevated levels of inflammatory checkpoints, including Nfkbia, Nfkbiz, Tnfaip3. In contrast, aged neutrophils had elevated expression of alarmins S100a6/a8/a9/a11 and other immune activation genes, including Retnlg, Lcn2, Ly6g, Cxcr2, Klf2, and Itgam (CD11b), largely matching the baseline trends, showing limited activation at this time point. The ISG signatures at this time point remained mixed, where young neutrophils had elevated levels of Isg15, Rsad2/Viperin, Oas3, Oasl2, Zc3hav1/ZAP, Parp14, Herc6, Ddx3x, while aged neutrophils had higher levels of Ifitm1/2/3/6, Oas1g, Ifi27, Slfn1, Slfn2, and Plscr1. Aged neutrophils also had elevated levels of MHC molecule H2-D1, H2- T22/T23/24, and H2-Q4/Q6/Q7. These data demonstrate that despite having lower baseline ISG signature, young neutrophils mount a strong antiviral response.

On day 7, neutrophils from aged mice had a clearly elevated interferon signature marked by increased expression of a large number of ISGs such as Ifitm3, Ly6e, Isg15, Ifit1/3/3b, Bst2, Irf7, Zbp1, Rtp4, Isg20, Ifi27l2a, Ifi203/209, and Ifi47. The aged neutrophils also had significantly elevated expression of inflammatory chemokines, including Ccl2, Ccl5, Cxcl10, and Cxcl2.

Expression of MHC-I molecules remained elevated at this time point in aged neutrophils, including H2-D1, Psmb8/10, and Psme2. On the other hand, young neutrophils on day 7 had elevated expression of immune-function related genes such as Itgal (CD11a), Csf3r, Entpd1 (CD39), Syk, Ptprc (CD45), Il31ra, Pde4d, and Notch2, along with chemotaxis and adhesion related genes (CD44, Dock2, and Vav3).

On day 11, aged neutrophils continued to express elevated ISG signatures, including Ifit3, Ifitm1/2, Irf1, Stat2, Ifi207, Ddx58, and Samhd1, indicating a lingering antiviral signature. Aged neutrophils also had significantly elevated levels of antigen presenting molecules including H2- K1, H2-Q7, H2-Q4, and H2-Q10 genes, indicating a persistent activation of neutrophils. At the same time, we observed a dramatic increase in inflammatory cytokines and chemokine gene expressions including Cxcl2, Cxcl3, Ccl2 (over log 5.5 fold change), and IL-1b in aged neutrophils. In contrast, young neutrophils on day 11 had elevated levels of cytokine receptor CXCR2, along with elevated levels of integrin genes Itgal and Itgax, genes associated with neutrophil functions, including Prkcb, Prkcd, Padi4 (encodes PAD4), Gsdme (Gasdermin E), Csf3r, MAP3K5, and Mmp9, indicating elevated functional status of young neutrophils on day 11.

These findings were supported by measurements of Influenza Signature Score (ISS), a representation of antiviral response [14], which peaked on day 3 in young mice and on day 7 in aged mice, indicating a delayed antiviral response in aged neutrophils (Fig. 3B). We then measured overall inflammatory response in neutrophils using the Inflammatory Response Score (IRS). Aged neutrophils had lower IRS at baseline and on day 3 but had an elevated score on day 7 and 11 post-infection (Fig. 3C).

Consistent with these gene expression data, our pathway analysis demonstrated that significantly enriched pathways in aged neutrophils on day 7 include “Chemokine receptors bind chemokines” and “Packaging of Eight RNA Segments”, while in young neutrophils, the enriched pathways include “IL-6-type cytokine receptor ligand interactions”, and “FCGR activation” (Fig. 3D). This trend continued at day 11 post-infection, where the most significantly upregulated pathways in aged neutrophils included “Interferon alpha/beta signaling” and “Antigen processing-Cross presentation” (Fig. 3E).

Together, these data indicate that aged neutrophils had dampened early antiviral and inflammatory responses during early influenza infection, followed by persistent antiviral and inflammatory responses.

### Neutrophils subsets demonstrate age-dependent changes in influenza-infected lungs

Neutrophil heterogeneity is an emerging area of investigation [30]. Our data show that neutrophil population consisted of three unique subpopulations (Sup Fig. 2A), including cluster 4 (with high expression of Retnlg, S100a8, S100a9, Hp, and Csf3r, Mmp8), cluster 6 (Fosb, Ptgs2, Il1bos, Slc7a11, Cass4, and Il1r2), and cluster 20 (Ccl4, Cxcl2, Acod1, Ccrl2, Ccl3, Cled4d, and Il1b) (Sup Fig. 2B and 2C). Pseudotime analysis revealed population 20 as the final stage of neutrophil trajectory (Sup Fig. 2D). The neutrophil heterogeneity was confirmed in an independent cohort of young and aged female mice from a shorter time course from prior published study[31] (Sup Fig. 2E and F). Among the neutrophil subpopulations, neutrophil 20 had higher ISS and IRS (Sup Fig. 2 G-J), a phenotype also observed in the validation cohort (Sup Fig. 3).

To identify specific subpopulations contributing to exacerbated immune responses in aged mice, we investigated their DEG signatures. We separated population 20 (high antiviral/inflammatory) from populations 4 and 6 (low antiviral/inflammatory). A heatmap of top 20 genes at each time point is shown in Sup Fig. 4A and D. Our data show that aged neutrophil populations 4 and 6, but not population 20, demonstrated a dramatic increase in ISS and IRS scores on day 7 and remained elevated on day 11 despite lower scores on day 3, compared to young neutrophil counterparts (Sup Fig. 4B, C, E and F). These data demonstrate that low inflammatory subpopulation of neutrophils drive persistent antiviral and inflammatory signaling in aged mice.

### Macrophages from aged mice show early impaired but prolonged inflammation and antiviral signaling during the course of influenza infection

Alveolar macrophages (AMs), being the first line of immune-mediated defense in the lung, act as an important antiviral defense [32, 33]. AMs had differential gene expression profiles between young and aged mice during the course of influenza infection (Fig. 4A).

**Fig. 4.**
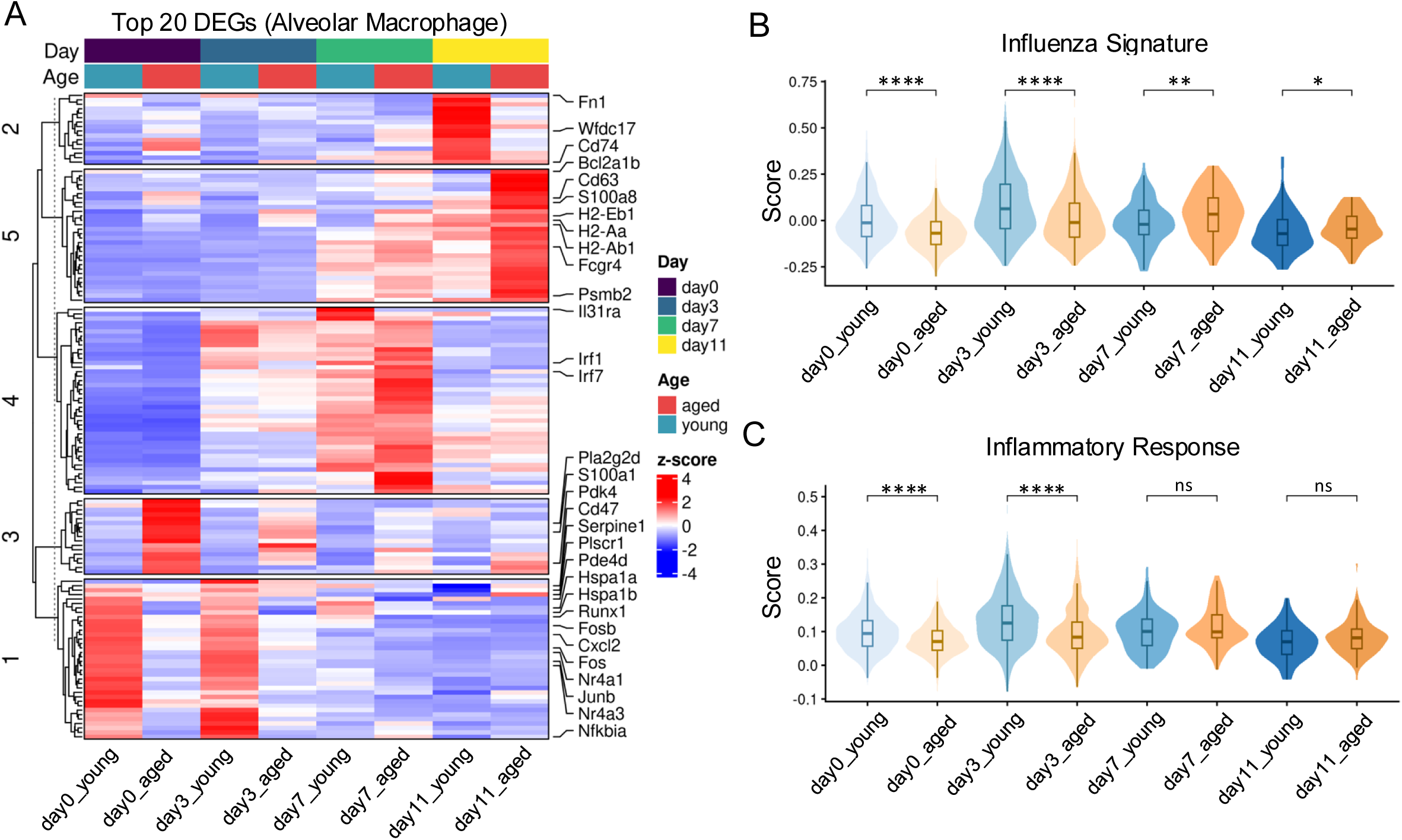
Alveolar macrophages demonstrate impaired early antiviral response but prolonged persistence of antiviral responses. Heatmap showing top 20 DEGs expressed at each time point between young and aged mice in alveolar macrophages (A). Quantification of ISS (B) and IRS (C) comparing young and aged AMs during the course of influenza infection. Data were analyzed using the Wilcoxon rank-sum test. *, P < 0.05; **, P < 0.01; ****, P < 0.0001; ns, not significant.

At baseline, aged AMs had elevated expression of MHC-II class molecules including H2-Aa, H2-Eb1, H2-Ab1, H2-K1, H2-DMb1, H2-M2, B2m, and H2-T22, (Fig. 4A, cluster 5), and genes related to immune activation and inflammation including CD63 (which supports interaction with P-selectin), CD74 (MIF receptor), S100a1/6/8/11, Ccl6, and Cxcr1. The most significantly upregulated gene in aged AMs was Pla2g2d, a putative anti-inflammatory phospholipase that impairs antiviral immunity [34]. In contrast, young AMs had elevated genes that are associated with more balanced inflammatory response including Nfkb1, Nfkbiz, Tnfa, Il6, Cxcl2, Nr4a1/3, Fos, Fosb, Fosl1/l2, Junb, Fn1 and Nfkbia (cluster 1 of the heatmap).

Pla2g2d remained elevated in aged AMs on day 3, along with multiple genes in MHC-II class H- 2 molecules. However, compared to aged AMs, young AMs demonstrated a significant increase in the HSP70 genes including both Hspa1a and Hspa1b (> 5 log fold change), along with antiviral genes (Irf1) and immunomodulator gene Mertk. On day 7 post-infection, there were fewer AMs in both young and aged mice, given “alveolar macrophage disappearance reaction” that depletes over 90% of AMs during influenza infection [35]. However, we observed that aged AMs had significantly elevated expression of H2-Ab1, Cd74, H2-Eb1, H2-Aa, while young AMs at this time point had higher expression of Pde4d and Runx1.

On day 11, aged AMs had increased expression of antiviral gene signature such as Irf7, Plscr1, Serpine1, Bcl2a1b, and Psmb2, a likely consequence of extended viral presence. Similarly, a significant increase in the pro-inflammatory genes was observed in AMs from aged mice including H2-K1 along with Fcgr4, Cd63, Cd47, Pdk4, and Ear genes (Ear1/2/6). In contrast, young AMs at this time point had elevated levels of immunomodulatory/reparative genes including Il31ra, Fn1, and Wfdc17, suggesting resolution of inflammation in young lungs at day 11.

Consistent with these gene changes, we observed that aged AMs had enrichment of “Interferon Signaling” and “MAPK1/MAPK3 signaling” pathways on day 11 post-infection. In contrast, young AMs had an enrichment in “Interleukin-4 and Interleukin-13 signaling” and “Extracellular matrix organization” pathways on day 11 (Sup Fig. 5A).

Aged AMs had decreased ISS at day 0 and day 3, indicating an impaired early antiviral ability. However, by day 7, AMs from aged mice had significantly higher ISS, a trend that continued on day 11, indicating elevated and persistent antiviral signature (Fig. 4B). Similar to the antiviral responses, IRS score demonstrated that aged mice had lower inflammatory response at day 0 and 3, however, at later stages of infection, aged AMs showed higher IRS (Fig. 4C). Combined with elevated viral burden in AMs from aged mice, our data indicate that aged AMs have impaired early sensing and antiviral response, resulting in persistent antiviral and inflammatory responses.

We observed similar findings in interstitial macrophages (IMs) and monocytes, which showed early impaired but a persistently elevated inflammatory and antiviral responses during the later course of infection (Sup Figures 5B, 6 and 7).

### Infection status of a cell is a major driver of transcriptional activity in myeloid cells

Our data show that aged mice have more infected cells compared to young mice, including multiple myeloid cells (Fig. 2D-G). Prior studies failed to account for the infection status of the cell and its impact on cellular transcriptional activity. We compared expression profiles of infected and uninfected cells from the same mice at the same time points to identify the effects of viral presence. We specifically focused on the aged group since they had significantly higher number of infected cells on day 11 post-infection. We confirmed that influenza genes are selectively expressed in infected cells (Fig. 5A).

**Fig. 5.**
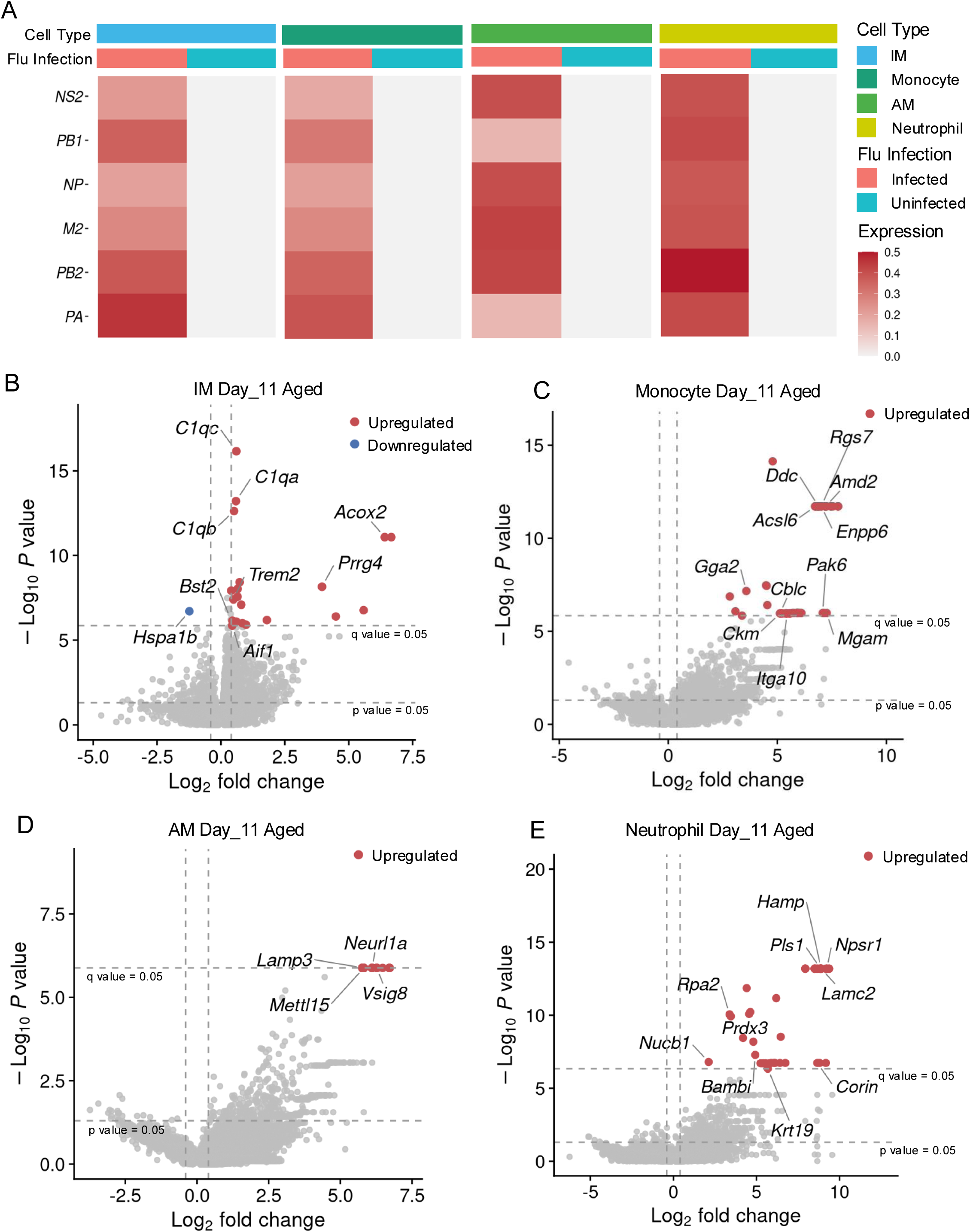
Infection status is a major driver of transcriptional activity in myeloid cells. Heatmaps demonstrating the expression of influenza genes in monocytes, IMs, AMs, and neutrophils, confirming selective expression of influenza genes in infected cells (A). Volcano plots demonstrating differentially expressed genes in IMs (B), monocytes (C), AMs (D), and neutrophils (E) obtained on day 11 post-infection from aged mice.

In aged IMs on day 11, we observed that infected cells had significantly higher expression of Bst2 (tetherin), a classical ISG, along with complement pathway genes (C1qc, C1qa, C1qb), MHC molecule H2-K1, and lysosomal enzyme Ctsz, suggesting an elevated inflammatory/pathological phenotype. At the same time, infected IMs had a significant decrease in the protective heat-shock protein 70 (Hspa1b), indicating lower host protective mechanisms in infected cells. Other elevated genes in infected cells vs uninfected cells include Acox2 (log2 fold change 6.4) and Prrg4 (log2 fold change 3.9), although their functional roles during influenza infection remain unclear (Fig. 5B).

Similar observations were made in AMs, neutrophils, and monocytes, where influenza infection was a major driver of transcriptional activity in these cells (Fig. 5C-E). Further, influenza-induced transcriptional upregulation was not limited to aged mice or day 11 post-infection. On day 3, we observed a significant upregulation of gene expression in infected vs uninfected IMs in both young (Sup Fig. 8A) and aged cells (Sup Fig. 8B). The elevated genes include Ifnb1, Il1a, and Il1rn in young and Cd40, Cxcl9, and Il1rn in aged IMs. These data suggest that influenza infection of immune cells is a major driver of transcriptional activation, regardless of age and disease stage.

### Aged endothelial cells have blunted early immune response but have persistent inflammatory responses during influenza infection

Endothelial cells form a critical barrier between lung and circulation, preventing fluid leak in the airspaces. We investigated how aging affects endothelial transcriptomic responses during influenza infection. Aged endothelial cells show “inflammaging phenotype” at baseline as evident by increased expression of genes including ISGs (Irf7, Isg15, Ifit1/3, Oas2/Oasl2, and Rsad2), antigen presentation genes (Cd74, H2-Ab1, and H2-Eb1), and inflammatory genes (Socs3, Cebpd, and Nfkbia). During early infection on day 3, young endothelial cells mounted robust antiviral and inflammatory responses compared to aged endothelial cells including higher expression of Nfkb1, Cxcl12, Tnfsf10, and Tnfrsf1a along with ISGs including Ifitm2/3, Ifi44/203/208/213, and Bst2 genes. At the same time, young endothelial cells expressed genes related to endothelial function and homeostasis (Cdh5, Pecam1, Tek, Tie1, Kdr, Flt1, Nrp1, and Egfl7), indicating a strong immune response while maintaining endothelial function (Fig. 6A).

**Fig. 6.**
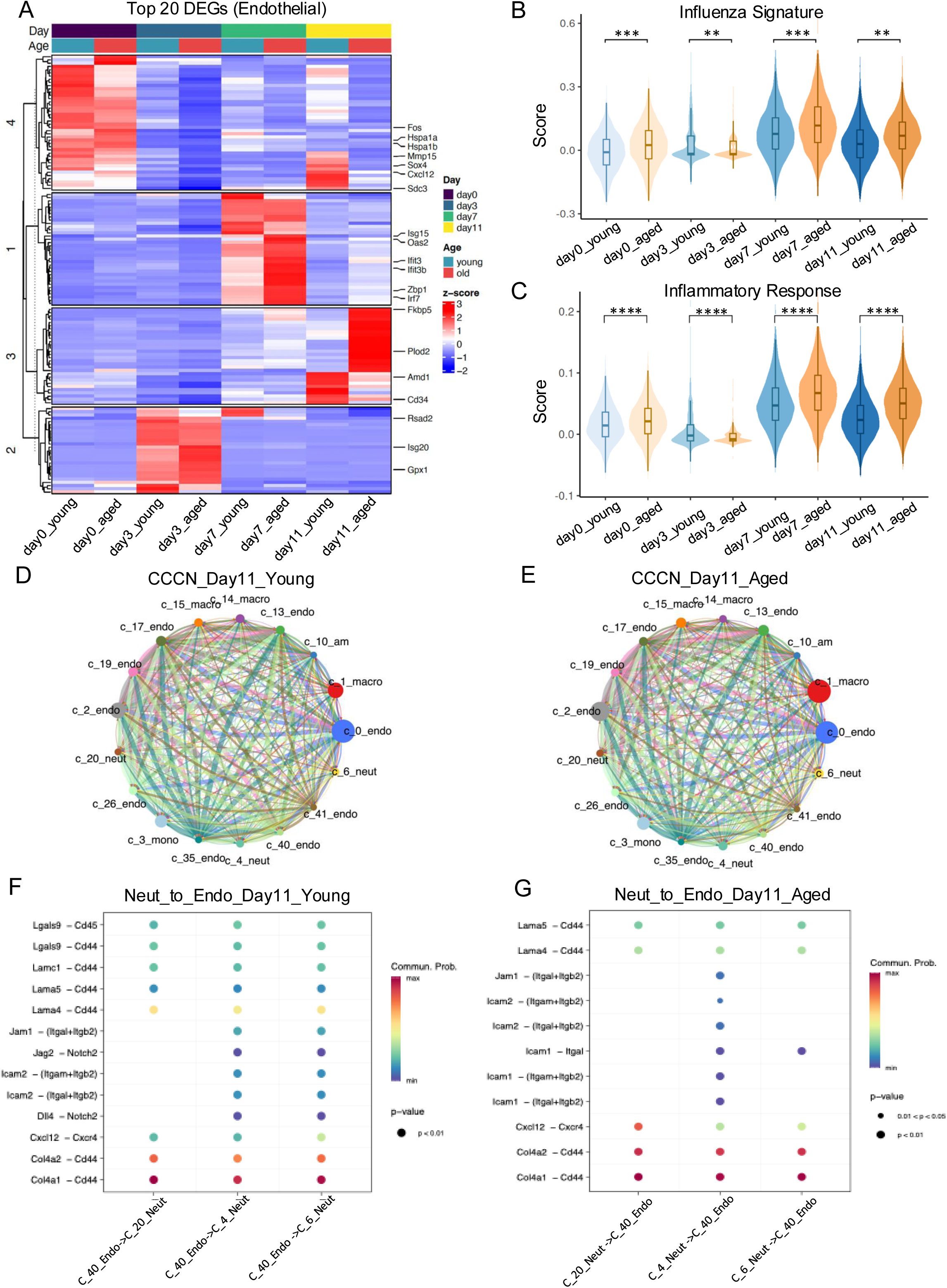
Endothelial cells demonstrate persistent antiviral and inflammatory responses during influenza infection. Heatmap showing top 20 DEGs expressed at each time point between young and aged mice in endothelial cells (A). IRS (B) and ISS (C) in young vs aged endothelial cells during the course of infection. Circos plots demonstrating cell-cell communication strengths between myeloid and endothelial cells in young (D) and aged (E) mice on day 11 post-infection. Specific ligand-receptor pair interactions on day 11 post-infection in young (F) and aged (G). Data were analyzed using the Wilcoxon rank-sum test. *, P < 0.05; **, P < 0.01; ****, P < 0.0001; ns, not significant.

The antiviral response in the aged endothelium was delayed to day 7 where numerous ISGs were elevated in aged endothelial cells including Ifit1/3, Isg15, Ifitm3, Oas2/3, Oasl2, Irf7, Gbp2/3/4/5, and Rsad2, among others. At this time point, endothelial cells in aged mice also had higher expression of adhesion molecules including Icam1/2 and Vcam1, a trend that continued on day 11 post-infection. A similar trend was observed in ISG signature where many ISGs remained elevated in aged endothelial cells on day 11. In contrast, young endothelial cells on day 11 expressed genes linked to endothelial identity and vascular remodeling, including Kdr, Aplnr, Pdgfb, Pdgfd, Prkg1, and Edil3 (Fig. 6A). These changes were reflected in IRS and ISS, which showed a persistent elevation in antiviral and inflammatory signatures in endothelial cells from aged mice on day 11 post-influenza infection (Fig. 6B and 6C).

Pathway analysis of endothelial cells demonstrated that on day 11 post-infection aged endothelial cells had enrichment of “Interferon alpha/beta signaling” and “p53-Dependent G1/S DNA damage checkpoint”, while the young endothelial cells showed enrichment of pathways including “Molecules associated with elastic fibers” and “RHO GTPase cycle”. A similar trend was observed on day 7 where a significant enrichment of “Viral RNP Complexes in the Host Cell Nucleus” and “Interferon gamma signaling” pathways was observed in aged endothelial cells, while young endothelial cells had a significant elevation of “IL-6-type cytokine receptor ligand interactions” and “Platelet homeostasis” pathways. These data indicate that aging renders the endothelial cells more inflammatory, likely contributing to the persistent inflammation and tissue injury (Sup Fig. 9).

### Aged lungs show altered interactions between endothelial and myeloid cells during the resolution phase of influenza infection

Endothelial-immune communications orchestrate inflammatory cell recruitment and resolution of inflammation. Aging altered cell-cell communications between multiple endothelial subsets and myeloid cells including neutrophils, alveolar macrophages, interstitial macrophages and monocytes (Fig. 6D, E and Sup Fig. 10 and 11).

On day 11, interactions between Col4a2 and CD44 were observed among neutrophil subsets and endothelial cells in both young and aged mice; however, these interactions were stronger in aged mice (Fig. 6F and G). Similarly, we observed that Cxcl12-Cxcr4 interactions were higher between aged endothelial cells and neutrophils, indicating a persistent inflammatory phase in the aged lungs. Other key differences include higher Lama4-Cd44 interactions in young while aged mice had higher Lama5-Cd44 interactions. In the reverse direction, endothelial to neutrophil (ligand to receptor) interactions were also affected by the aging process. These changes include a stronger interaction of oncostatin (Osm) with its receptors (Osmr+Il6st) and (Lifr+Il6st) in aging lung (Sup Fig. 10). Similar results were obtained on day 7 where we observed that aged endothelial cells had unique cell-cell communications with neutrophils such as Lgals9-Cd44 and Lgals9-Cd45 (Sup Fig. 11). Taken together, these cell-cell communications indicate underlying molecular mechanisms that drive persistent lung injury in the aged host.

### Sex differences are present in anti-influenza responses across multiple cell types

Influenza infections show sex bias where young females are more susceptible to infection compared to young males [3]; however, mortality largely occurs in the aged males [5]. We investigated the sex-dependent transcriptomic differences in the immune cells during influenza infection. Comparisons were made within the same time point and age group to identify sex-specific differences. We validated our analytical approach by examining Y chromosome (Ddx3y, Uty, and Eif2s3y) and X chromosome genes (Xist, Tsix and Kdm6a) (Sup Fig. 12).

In young neutrophils at baseline, females had higher expression of ISGs such as Ifitm2 and Slfn2, along with the inflammatory genes such as Il1b, Tnfaip2, Nfkbid, and Nfkbia, and C5ar1, and alarmins including S100a6/8/9/11 (Fig. 7A). Young male neutrophils at this time point had elevation of inflammatory genes such as Pf4 (Cxcl4), Ppbp (Cxcl7), Osm (Oncostatin M), Il15, Ccr1, Map3k5, and Cebpb (Fig. 7A). Interestingly, aged male neutrophils had higher expression of many of inflammatory genes compared to aged female neutrophils including Ccl3, Ccl4, Cxcl2, Nfkb1, Nfkbiz, Nfkbie, Tnfaip3 (A20), and Nr4a3 (Fig. 7B), indicating higher inflammatory tone in aged male neutrophils at baseline. Aged female neutrophils had higher expression of S100 genes including S100a6/8/9/11, Ccl6, Ltb, Mmp9, Alox5ap, Pglyrp1, and Lyz2.

**Fig. 7.**
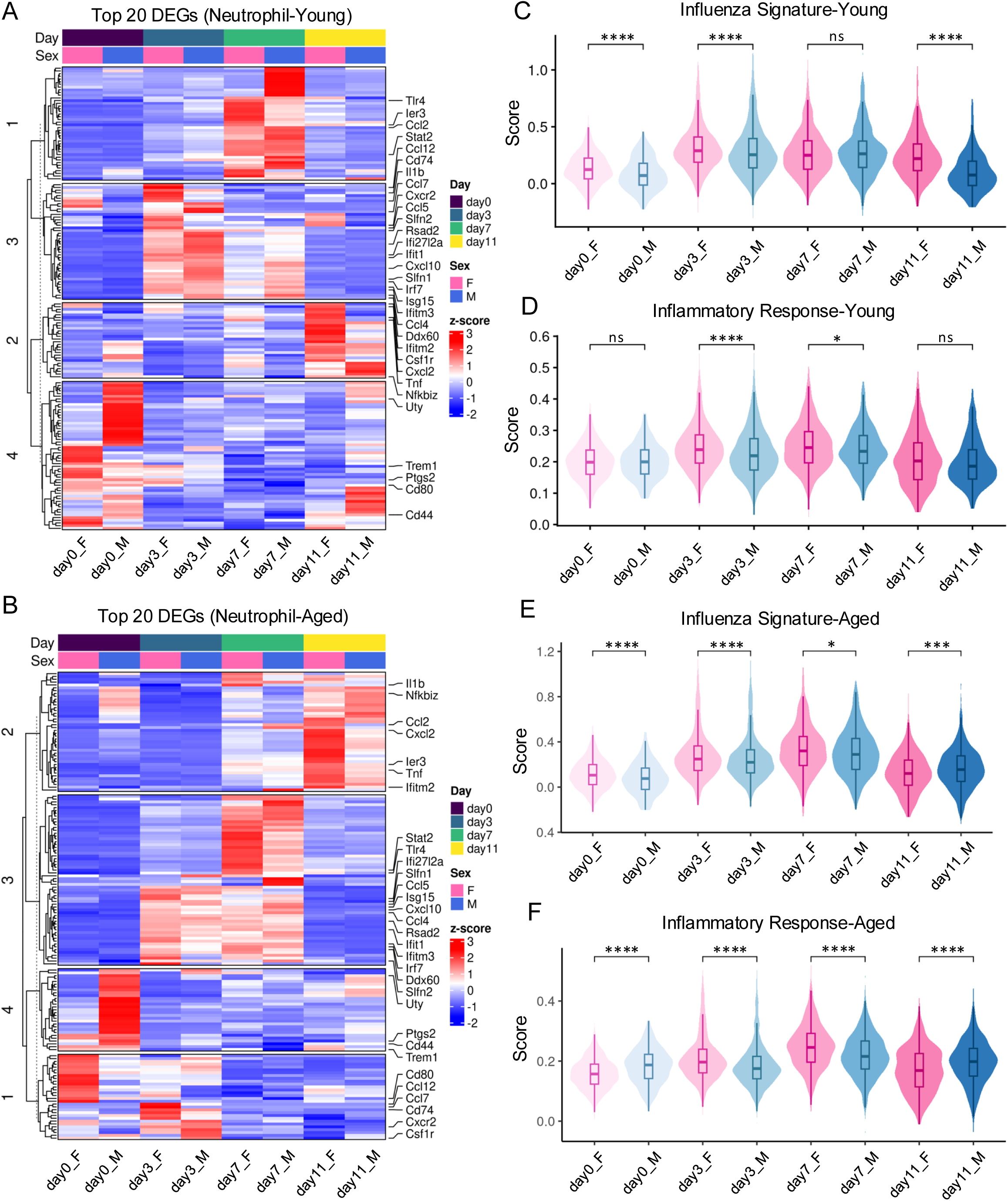
Sex differences in neutrophil gene expression in young and aged mice. Heatmap showing the top 20 DEGs expressed at each time point between male and female neutrophils during the course of influenza infection in young (A) and aged (B) mice. ISS and IRS differences between male vs female cells in young (C and D) and aged (E and F) mice. Data were analyzed using the Wilcoxon rank-sum test. *, P < 0.05; **, P < 0.01; ***, P < 0.001; ns, not significant.

On day 3 post-infection, among the young, we observed that female neutrophils had increased expression of numerous inflammatory genes including Il1b, Cxcl2, Cxcr2, Ptgs2 (COX-2), Selplg (PSGL-1), Nfkbid, Nfkbiz, Cd14, Mmp9, alarmin genes (S100a8/a9/a11), and antiviral genes such as Ifitm1/2/3/6. Similarly, aged female neutrophils had a higher interferon response manifested by elevated expression of multiple ISGs, including Ifit1/2/3/3b, Oas1a, Oas1g, Oasl2, Ifi27l2a, Ifi204, Ifitm3, Isg15/20, Xaf1, Zbp1, Rtp4, Ddx6/60, Stat2, and Ube2l6. Aged female neutrophils also had higher expression of many inflammatory genes including Nfkbiz, Ier3, Tlr4, C5ar1, Tarm1, Padi4, Ly6g, Ly6c2, Cd177, Mmp8/25, H2-K1/H2-Q4/-Q6/-Q7, while aged males had elevated expression of Myadm, Bpifa1, and Il1b.

On day 7 post-infection, young female neutrophils had higher expression of Clec4d, H2-Q6, H2- Q7, Cd44, Alcam, Ctsb, and Nfkbid genes while male neutrophils had higher expression of ISGs including Isg15, Ifit1, Ifit3, and Rsad2 (viperin) (Fig. 7A). In the aged mice on day 7, we observed that females had significantly higher expression of ISGs compared to males, a pattern opposite to that seen in young neutrophils on day 7. The elevated ISGs in females include Gbp2, Gbp7, Oasl2, Ly6e, Irf1, and Igtp, while a subset of ISGs including Isg15 and Ifi27l2a remained elevated in aged male neutrophils. Aged female mice also had higher expression of inflammatory genes including Csrnp1, Ly6a, Ly6e, Cd44, Nfkbiz, and Zfp36 on day 7 (Fig. 7B).

On day 11 post-infection, in young female neutrophils, there was upregulation of ISGs including Ifitm1/2/3, although most other ISGs were not differentially expressed at this time point. Young female neutrophils also had elevated expression of alarmin-mediated inflammatory signature including S100a8/9, Cd14, and Wfdc17. Young male cells at day 11 had elevated expression of MAPK genes (Mapk14 and Map3k5) and Dusp1, a negative regulator of MAPK. Other significantly elevated genes in young male neutrophils include Il1r2 (decoy receptor for IL-1), Txnip, Fos, Fosb, Junb, Jund, and Gsr, indicating strong anti-inflammatory and resolution phenotype in male neutrophils (Fig. 7A). In contrast, aged male neutrophils on day 11 had elevated expression of numerous inflammatory genes including Pde4b, Ptprc, Csf3r, Sell, Cd52, Mmp9, Arg2, Ptgs2 (COX-2), Tnfaip2, and Cyp4f18. Aged female neutrophils had higher expression of chemokines including Cxcl2/3, Ccl2, Cebpb, and Cd14 genes and oxidative stress related genes such as Hilpda, Txnip, and Mt1. Most of the classical ISGs at this time point were not statistically different between males and females.

We also investigated ISS and IRS between male and female neutrophils in both young and aged mice (Fig. 7C-F). Overall, females had higher ISS and IRS throughout infection, consistent with stronger antiviral and inflammatory responses (Fig. 7C and D). However, aged males had higher ISS at baseline and higher ISS and IRS on day 11 post-infection indicating that aged males had persistent inflammatory and antiviral responses post-influenza infection (Fig. 7E and F). This pattern was consistent across multiple myeloid compartments including AMs, IMs and monocytes (Sup Figs. 13, 14, and 15). Taken together, these data show that while young male myeloid cells generally downregulate inflammatory and antiviral programs during the resolution phase, aged male myeloid cells maintain heightened ISS and IRS signatures, revealing persistent immune activation in aged males.

### Aging induced immune responses in human lung macrophage populations during COVID- 19 infection

To investigate whether aged myeloid cells show similar age-dependent phenotypes in humans, we analyzed publicly available single-cell RNA sequencing data from young and aged COVID- 19 patients and healthy individuals. Among the alveolar macrophages from COVID-19 patients, we observed a clear dichotomy in differentially expressed genes between young and aged individuals. AMs from aged COVID-19 patients showed a clear elevation of numerous ISGs including MX1, IFITM1/2/3, IFI30/35, IFIT2/3, ISG15, PYCARD, S100A13, BST2, OASL, IRF7, STAT1, and OAS3, among others. On the other hand, young cells showed upregulation of genes related to repair/inflammatory regulation including AREG, FOS/B, JUND, TIMP2, VEGFB, FOSL2, along with immune genes such as CXCR4 and MAPK1, indicating a balanced inflammatory response (Fig. 8A). Consistent with these gene expression changes, we observed that aged AMs had enrichment of pathways including “Cytokine Signaling in Immune system”, and “Interferon alpha/beta signaling”, while the enriched pathways in young AMs post-infection included “L13a-mediated translational silencing of Ceruloplasmin expression” and “GTP hydrolysis and joining of the 60S ribosomal subunit”.

**Fig. 8.**
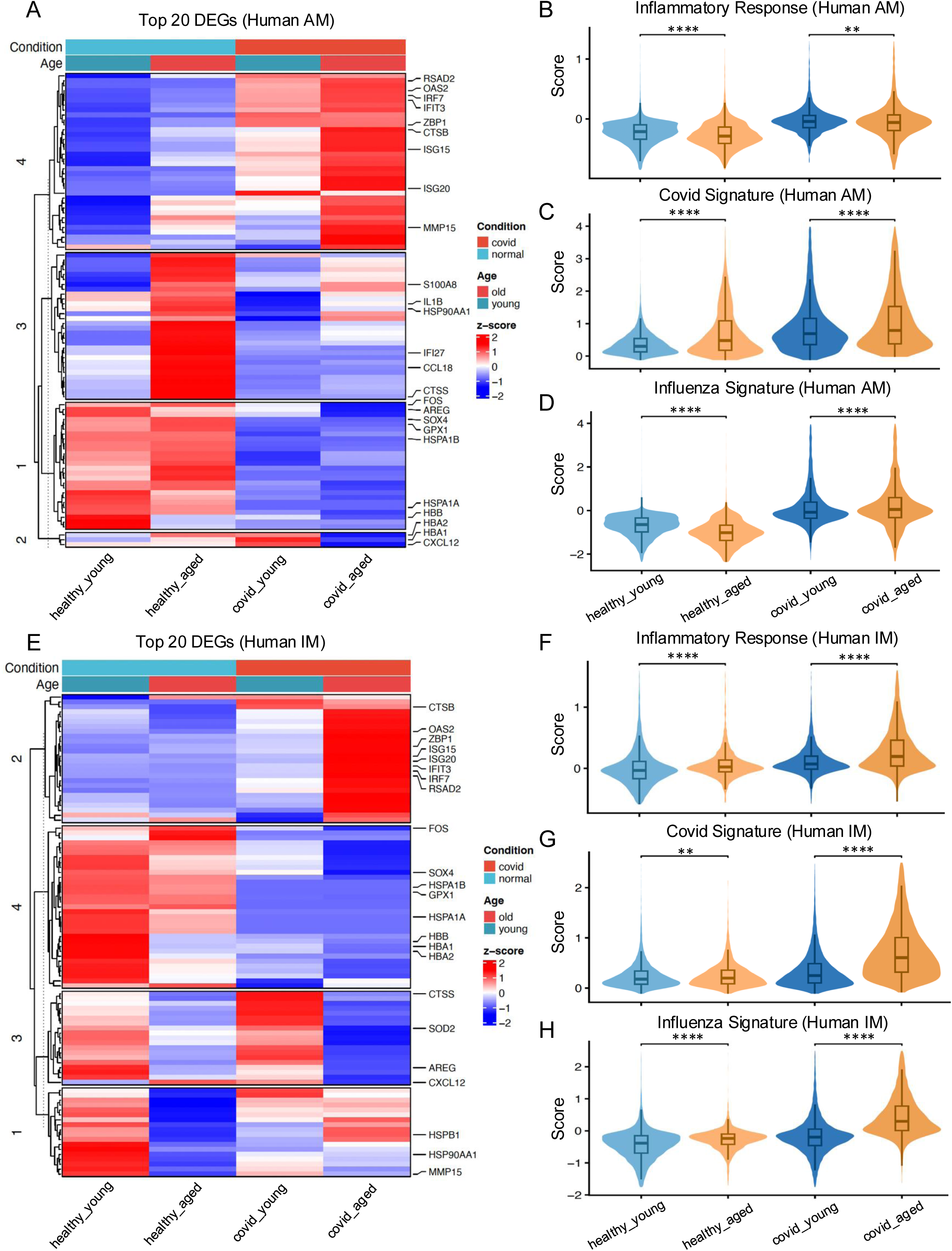
Aged human macrophages show elevated antiviral and inflammatory gene signatures during COVID-19. A heatmap showing the top 20 DEGs expressed in healthy or COVID-19 immune cells in alveolar macrophages (A) and interstitial macrophages (B). ISS, IRS, and CRS in Alveolar macrophages (C-E) and interstitial macrophages (F-H). Data were analyzed using the Wilcoxon rank-sum test. *, P < 0.05; **, P < 0.01; ***, P < 0.001; ns, not significant.

We also quantified overall inflammatory (IRS) and antiviral (ISS) responses, along with a COVID-19 Response Score (CRS) in young and aged human AMs from healthy individuals and COVID-19 patients. Our data show that among the healthy individuals, aged AMs had comparable or lower IRS, ISS, but elevated CRS scores (Fig. 8D-F). However, post-infection all three scores were higher in aged AMs, indicating that aging leads to elevated inflammatory and antiviral gene signature in aged individuals (Fig. 8D-F), consistent with mouse findings at day 7 and day 11 post-infection.

Similar findings were observed in interstitial macrophages and monocytes from aged humans during COVID-19, which demonstrated higher expression of inflammatory and antiviral immune activation markers compared to young patients with COVID-19 (Fig. 8E- G, and Sup Fig. 16).

Taken together, these data indicate that aging alters antiviral and inflammatory responses in both mice and humans, contributing to persistent lung inflammation.

## Discussion

Aging is a critical determinant of viral infection severity, yet the cellular and molecular mechanisms underlying age-related susceptibility remain incompletely understood. By integrating single-cell transcriptomics across age, sex, and multiple infection stages in mice and human BAL samples from COVID-19 patients and healthy individuals, our study reveals previously unrecognized determinants of antiviral immunity and disease severity in aged hosts. A key conceptual advance is that the transcriptional landscape of the infected lung is shaped not only by age and sex, but also profoundly by the infection status of individuals cells. Myeloid cells containing viral transcripts exhibited widespread and coordinated upregulation of interferon-stimulated genes, antigen-presentation pathways, complement components, cathepsins, oxidative stress programs, and alarmins. Given that aged mice harbor a substantially greater proportion of infected myeloid cells during the late phase of disease (Fig. 2), this infection status-driven activation results in prolonged inflammation and delayed resolution.

Across neutrophils, AMs, IMs, and monocytes, aging resulted in a pattern of immune dysregulation: blunted early antiviral responses, followed by elevated and persistent inflammatory activation at later stages of the disease [36]. This delayed and persistent inflammatory response is linked to the prolonged presence of viral RNA in aged lungs, suggesting that ineffective early viral control impairs tissue repair, prolonging lung injury. Notably, low-inflammatory subsets of neutrophils contribute to low-grade persistent inflammation and extended antiviral responses in aged mice. This indicates that aging not only alters the magnitude of immune activation but also changes the functional roles of specific cellular subsets.

Importantly, despite the absence of live pathogens at later stages of the disease, viral pathogen components are sufficient to contribute to persistent inflammatory and injury responses. This concept has been supported by prior studies showing viral remnants contribute to the host response months after viral clearance [37]. Thus, our data provide unique insight into influenza pathology and a potential therapeutic target for limiting prolonged disease independently of active viral replication.

Another key insight from our analysis is the profound interaction between age and sex. While young females displayed stronger early antiviral (ISS) and inflammatory (IRS) signatures, consistent with known female-biased antiviral immunity, this pattern reversed with aging. Aged males exhibited elevated late-phase inflammatory and antiviral signaling across all major myeloid compartments. Importantly, aged male myeloid cells maintained elevated transcriptional activation even when viral RNA levels began to decline, suggesting intrinsic dysregulation in the resolution phase of lung injury. This persistent inflammatory state parallels clinical observations that aged men experience disproportionately higher hospitalization and mortality rates from influenza and other respiratory viral infections [6]. Our findings provide a mechanistic foundation for these epidemiological trends, indicating that age-dependent remodeling of sex-specific immunity, rather than sex alone, determines susceptibility to adverse outcomes.

In conclusion, this study provides a comprehensive single-cell resolution map of how age, sex, and infection status intersect to shape antiviral and inflammatory responses in the lung. Our findings reveal that viral persistence in aged myeloid cells and its interaction with endothelial cells drives a feedforward cycle of inflammation and impaired resolution, with aged males showing the most pronounced and prolonged immune activation. These age- and sex-specific transcriptional programs are conserved in human COVID-19, underscoring their translational relevance. Together, our atlas provides a foundational resource for understanding age- and sex-dependent vulnerability to respiratory viral infections and a framework for developing targeted therapeutic strategies in high-risk populations.

## Supporting information

All supplemental figures combined

## Limitations

Here we provide a comprehensive time-resolved single-cell RNA sequencing atlas of influenza-infected lung in young and aged mice of both sexes. However, there are certain limitations of this study worth acknowledging. First, while each time point included n=2 biological samples per sex (n=4 per age group), we acknowledge that higher biological replication would further strengthen the statistical power of age-by-sex interaction analyses, particularly for subtle transcriptional differences. Second, infected cells were defined by the expression of at least one viral transcript following DecontX correction to minimize ambient RNA contamination; however, we acknowledge that this threshold may include cells that phagocytosed viral debris rather than productively infected cells, and stricter thresholds may refine these findings in future studies. Third, mouse models do not recapitulate all aspects of human disease including sex differences. For example, while high susceptibility in young females is well replicated in mice, aged female mice remain more susceptible to influenza than aged males, a likely consequence of lower body weight in aged female mice when using the same inoculum dose. Fourth, cell-cell communication predictions between endothelial and immune cells are based on complementary receptor-ligand gene expression and require experimental validation to confirm these interactions in a spatial and functional manner. Finally, COVID-19 is a pathologically and mechanistically distinct disease compared to influenza infection; however, in the absence of a reliable human influenza cohort across the age spectrum, we utilized these publicly available data to identify conserved molecular signatures across common respiratory viral diseases.

## Supplementary figures

**Sup Fig. 1. Cellular proportion during the course of influenza in the lung.** Stacked plot showing proportion of cells during the course of influenza infection between young and aged mice.

**Sup Fig. 2. Neutrophil heterogeneity during influenza infection.** UMAP demonstrating three unique neutrophil subpopulations present in our data (A). The unique gene signatures that define each of the neutrophil subpopulations are shown (B). Heatmap demonstrating the unique gene signature of each of the three subpopulations (C). Pseudotime analysis demonstrating the trajectory of neutrophil maturation (D). Three distinct neutrophil subpopulations are also observed in the validation cohort (E), along with the gene signatures for each neutrophil subpopulation (F). UMAP showing the intensity of the Influenza Signature Score (ISS) (G) and its quantification (H). UMAP demonstrating the intensity of Inflammatory Response Score (IRS) (I) and its quantification (J) across three distinct neutrophil subpopulations. Data were analyzed using the Wilcoxon rank-sum test, ****, P < 0.0001.

**Sup Fig. 3. Validation of the differential ability of three neutrophil subpopulations to exert antiviral and inflammatory responses using independent publicly available dataset.** UMAPs showing Influenza Signature Score (A) and Inflammatory Response Scores (B) in the validation cohort. The quantification of ISS (C) and IRS (D) in different neutrophil subpopulations in the validation cohort. Data were analyzed using the Wilcoxon rank-sum test. *, P < 0.05; **, P < 0.01; ***, P < 0.001; ****, P < 0.0001.

**Sup Fig. 4. Neutrophils show cluster-specific changes between young and aged mice.** Heatmap demonstrating time-dependent gene expression changes between young and aged neutrophil cluster 20, generated using the top 20 DEGs in the group (A). Violin plots demonstrating ISS (B) and IRS (C) in neutrophil cluster 20. Heatmap demonstrating time-dependent gene expression changes between young and aged neutrophil 4 and 6 clusters generated using the top 20 DEGs in the group (D). Violin plots demonstrating ISS (E) and IRS (F) in neutrophil clusters 4 and 6. Data were analyzed using the Wilcoxon rank-sum test. *, P < 0.05; **, P < 0.01; ***, P < 0.001; ****, P < 0.0001; ns, not significant.

**Sup Fig. 5. Pathway enrichment in alveolar and interstitial macrophages on day 11 post-infection.** Upregulated pathways in aged mice (shown in red) and upregulated pathways in young (shown in blue) on day 11 in AMs (A) and in IMs (B).

**Sup Fig. 6. Aging alters antiviral and inflammatory responses in interstitial macrophages.** Heatmap showing top 20 DEGs expressed at each time point between young vs aged mice in Ims (A). Quantification of ISS (B) and IRS (C) comparing young and aged IMs during the course of influenza infection. Data were analyzed using the Wilcoxon rank-sum test. *, P < 0.05; ****, P < 0.0001.

**Sup Fig. 7. Aging alters antiviral and inflammatory responses in monocytes.** Heatmap demonstrating time-dependent gene expression changes of monocytes between young and aged mice generated using the top 20 DEGs in the group (A). Violin plots demonstrating ISS (B) and IRS (C) in monocytes comparing young and aged monocyte scores during the course of influenza infection. Data were analyzed using the Wilcoxon rank-sum test. *, P < 0.05; ***, P < 0.001; ****, P < 0.0001.

**Sup Fig. 8. Infection of IMs drives transcriptional activity in both young and aged mice on day 3 post-infection.** Volcano plots demonstrating upregulated genes in infected IMs compared to uninfected IMs obtained on day 3 post-infection from young (A) and aged (B) mice. The horizontal dashed lines denote p-value (lower) < 0.05 and q-value (upper) < 0.05.

**Sup Fig. 9. Pathway enrichment in endothelial cells on day 7 and day 11 post-infection.** Upregulated pathways in aged mice (shown in red) and upregulated pathways in young (shown in blue) endothelial cells shown on day 7 (A) and day 11 (B).

**Sup Fig. 10. Ligand-receptor interactions between endothelial and neutrophils on day 11 post-infection.** Cell-cell communication plots demonstrating interactions between endothelium (source of ligand) and neutrophils (source of receptor) on day 11 post-infection.

**Sup Fig. 11. Altered cell-cell interactions between endothelial and neutrophils on day 7 post-infection.** Circos plots demonstrating overall cell-cell interactions between young (A) and aged (B) mice on day 7 post-infection. Specific ligand-receptor pairs where neutrophil acts as a source of ligand and endothelial cells as a receptor between young (C) and aged (D) mice. Interaction strength between ligand (endothelial)-receptor (neutrophils) pairs between young (E) and aged (F) on day 7 post-infection.

**Sup Fig. 12. Sex linked genes show differential expression between male and female cells.** Gene expression of three X chromosome-linked genes (Xist, Tsix, and Kdm6a) and three Y chromosome-linked genes (Ddx3y, Uty, and Eif2s3y). The expression patterns are shown by the combination of cell types (IMs, monocytes, AMs, and neutrophils) and sex (female and male).

**Sup Fig. 13. AMs exhibit sex differences during the course of influenza infection.** Heatmap demonstrating time-dependent gene expression changes between male and female AMs generated using the top 20 DEGs in young (A) and aged mice (B). ISS and IRS differences between male vs female cells in young (C and D) and aged (E and F) mice. Data were analyzed using the Wilcoxon rank-sum test. *, P < 0.05; **, P < 0.01; ***, P < 0.001; ns, not significant.

**Sup Fig. 14. Interstitial macrophages exhibit sex differences during the course of influenza infection.** Heatmap demonstrating time-dependent gene expression changes between male and female IMs generated using the top 20 DEGs in young (A) and aged mice (B). ISS and IRS differences between male vs female cells in young (C and D) and aged (E and F) mice. Data were analyzed using the Wilcoxon rank-sum test. *, P < 0.05; **, P < 0.01; ***, P < 0.001; ns, not significant.

**Sup Fig. 15. Monocytes exhibit sex differences during the course of influenza infection.** Heatmap demonstrating time-dependent gene expression changes between male and female monocytes generated using the top 20 DEGs in young (A) and aged mice (B). ISS and IRS differences between male vs female cells in young (C and D) and aged (E and F) mice. Data were analyzed using the Wilcoxon rank-sum test. *, P < 0.05; **, P < 0.01; ***, P < 0.001; ns, not significant.

**Sup Fig. 16. Aging alters antiviral and inflammatory responses in human monocytes.** Heatmap demonstrating time-dependent gene expression changes of monocytes between young and aged human monocytes generated using the top 20 DEGs in the group (A). Violin plots demonstrating IRS (B), CRS (C), and ISS (D) in human monocytes comparing young and aged monocyte scores. Upregulated pathways in aged humans cells (shown in red) and upregulated pathways in young human cells (shown in blue) in AMs (E), IMs (F), and monocytes (G) from COVID-19 infected individuals. Data were analyzed using the Wilcoxon rank-sum test. ****, P < 0.0001.

