## Supplementary material for "A Time-Resolved Single-Cell Atlas Reveals Infection-Status, Age-, and Sex-Dependent Immune Responses Drive Viral Disease Severity": All supplemental figures combined

Sup Fig. 1

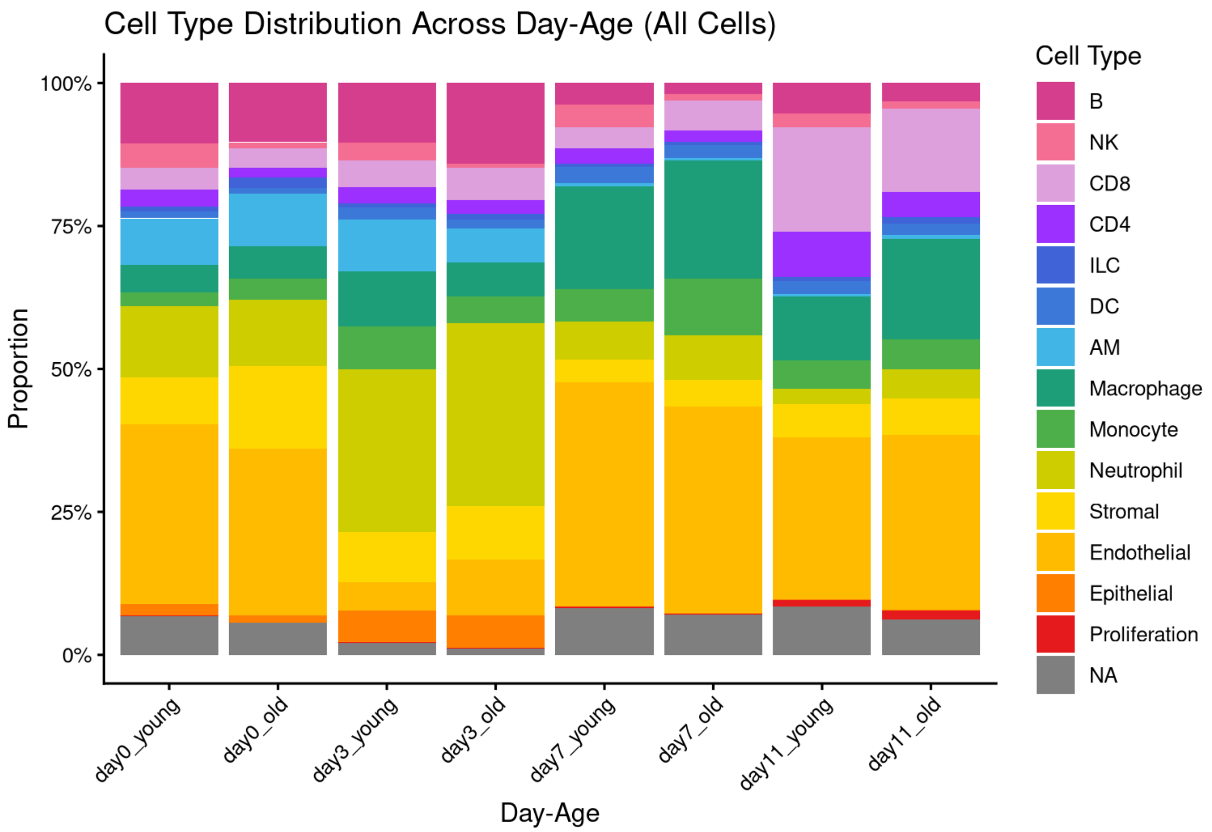

Cell Type Distribution Across Day-Age (All Cells)
Proportion
100%
75%
50%
25%
0%
day0_young
day0_old
day3_young
day3_old
day7_young
day7_old
day11_young
day11_old
Day-Age
Cell Type
B
NK
CD8
CD4
ILC
DC
AM
Macrophage
Monocyte
Neutrophil
Stromal
Endothelial
Epithelial
Proliferation
NA

Sup Fig. 2

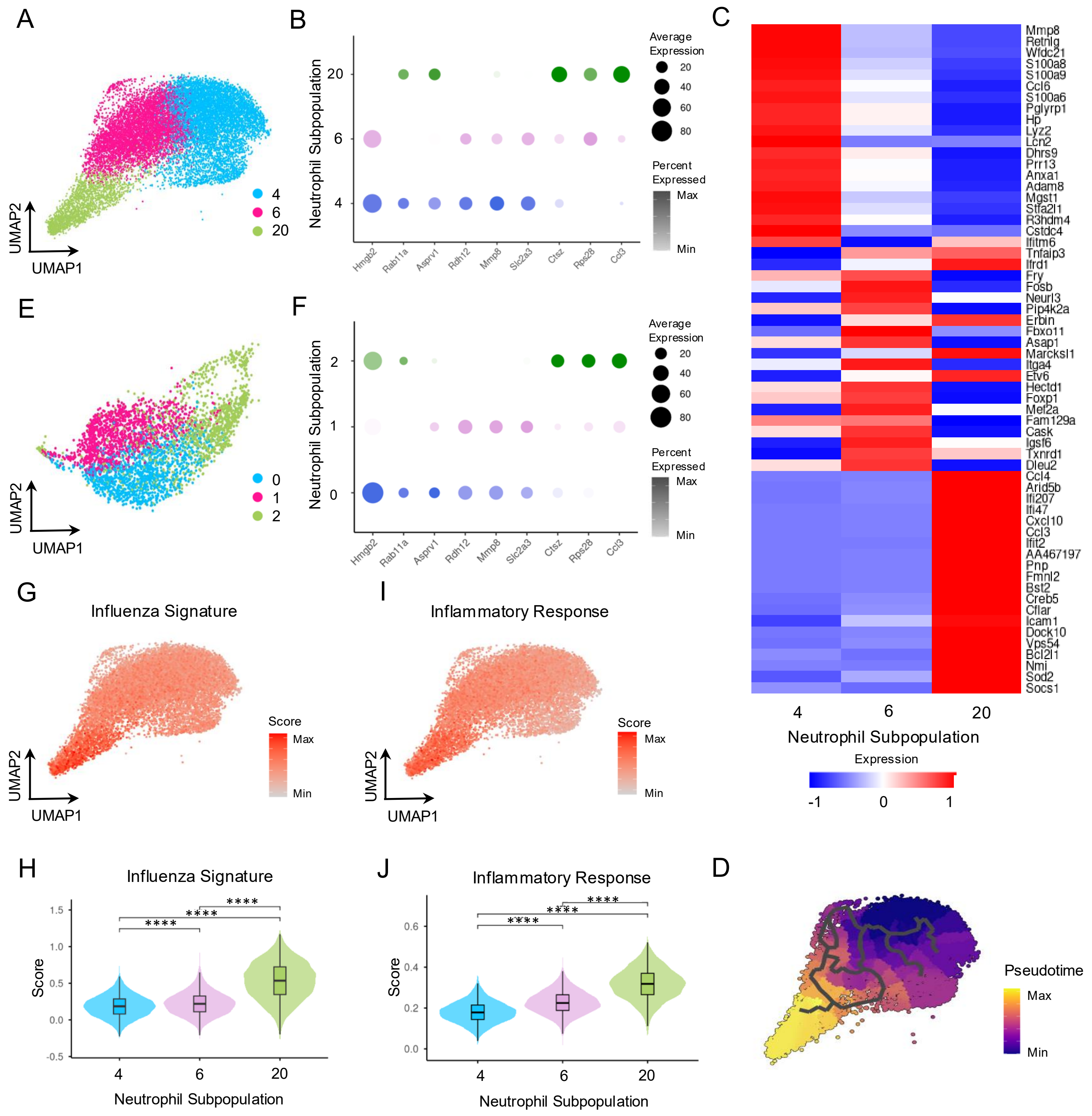

A
UMAP2
UMAP1
4
6
20
B
Neutrophil Subpopulation
Average Expression
Percent Expressed
Max
Min
Hmgb2
Rab11a
Asprv1
Rdh12
Mmp8
Slc2a3
Ctsz
Rps28
Ccl3
C
Mmp8
Retnlg
Wfdc21
S100a8
S100a9
Ccl6
S100a6
Pglyrp1
Hp
Lyz2
Lcn2
Dhrs9
Prr13
Anxa1
Adam8
Mgst1
Stfa2l1
R3hdm4
Cstdc4
Ifitm6
Tnfaip3
Ifrd1
Fry
Fosb
Neurl3
Pip4k2a
Erbin
Fbxo11
Asap1
Marcksl1
Itga4
Etv6
Hectd1
Foxp1
Mef2a
Fam129a
Cask
Igsf6
Txnrd1
Dleu2
Ccl4
Arid5b
Ifi207
Ifi47
Cxcl10
Ccl3
Ifit2
AA467197
Pnp
Fmnl2
Bst2
Creb5
Cflar
Icam1
Dock10
Vps54
Bcl2l1
Nmi
Sod2
Socs1
Neutrophil Subpopulation
Expression
E
F
G
Influenza Signature
Score
I
Inflammatory Response
H
Influenza Signature
Score
Neutrophil Subpopulation
J
Inflammatory Response
D
Pseudotime

Sup Fig. 3

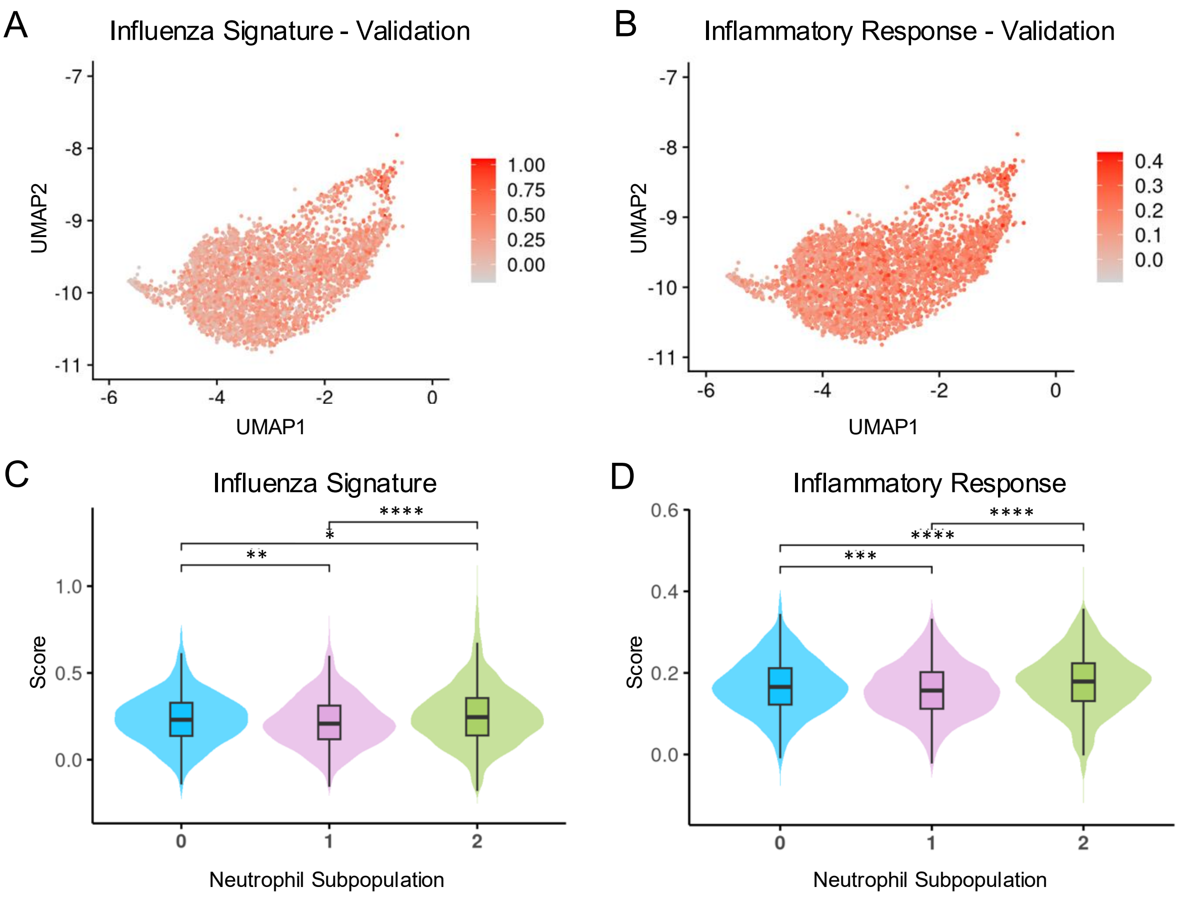
A
Influenza Signature - Validation
UMAP2
UMAP1
-7
-8
-9
-10
-11
-6
-4
-2
0
1.00
0.75
0.50
0.25
0.00
B
Inflammatory Response - Validation
UMAP2
UMAP1
-7
-8
-9
-10
-11
-6
-4
-2
0
0.4
0.3
0.2
0.1
0.0
C
Influenza Signature
Score
Neutrophil Subpopulation
1.0
0.5
0.0
0
1
2
****
*
**
D
Inflammatory Response
Score
Neutrophil Subpopulation
0.6
0.4
0.2
0.0
0
1
2
****
****
***

Sup Fig. 4

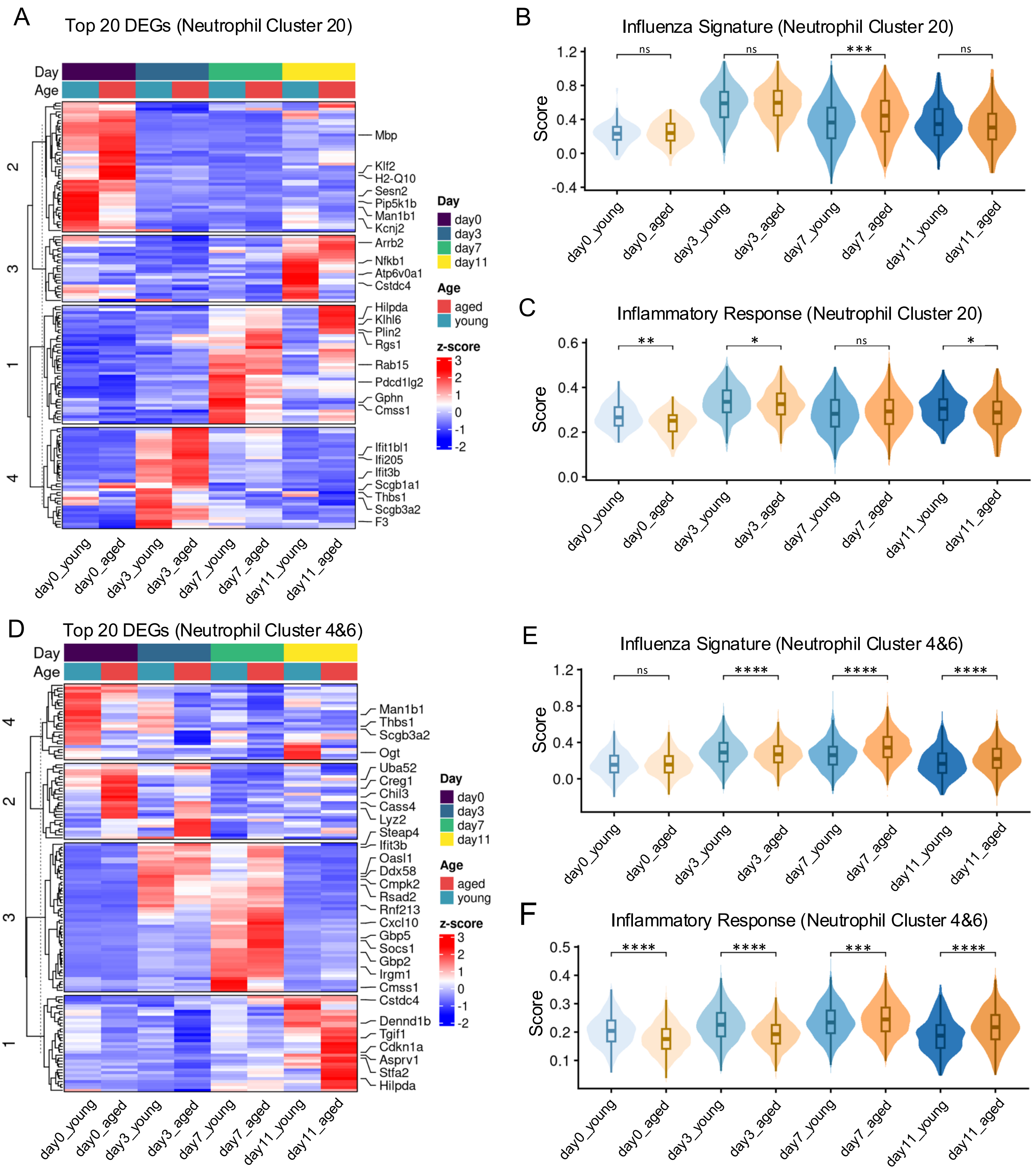
A
Top 20 DEGs (Neutrophil Cluster 20)
Day
Age
Mbp
Klf2
H2-Q10
Sesn2
Pip5k1b
Man1b1
Kcnj2
Arrb2
Nfkb1
Atp6v0a1
Cstdc4
Hilpda
Klhl6
Plin2
Rgs1
Rab15
Pdcd1lg2
Gphn
Cmss1
Ifit1bl1
Ifi205
Ifit3b
Scgb1a1
Thbs1
Scgb3a2
F3
day0_young
day0_aged
day3_young
day3_aged
day7_young
day7_aged
day11_young
day11_aged
Day
day0
day3
day7
day11
Age
aged
young
z-score
B
Influenza Signature (Neutrophil Cluster 20)
Score
ns
ns
***
ns
C
Inflammatory Response (Neutrophil Cluster 20)
Score
**
*
ns
*
D
Top 20 DEGs (Neutrophil Cluster 4&6)
Man1b1
Thbs1
Scgb3a2
Ogt
Uba52
Creg1
Chil3
Cass4
Lyz2
Steap4
Ifit3b
Oasl1
Ddx58
Cmpk2
Rsad2
Rnf213
Cxcl10
Gbp5
Socs1
Gbp2
Irgm1
Cmss1
Cstdc4
Dennd1b
Tgif1
Cdkn1a
Asprv1
Stfa2
Hilpda
E
Influenza Signature (Neutrophil Cluster 4&6)
ns
****
****
****
F
Inflammatory Response (Neutrophil Cluster 4&6)
****
****
***
****

### Sup Fig. 5

A

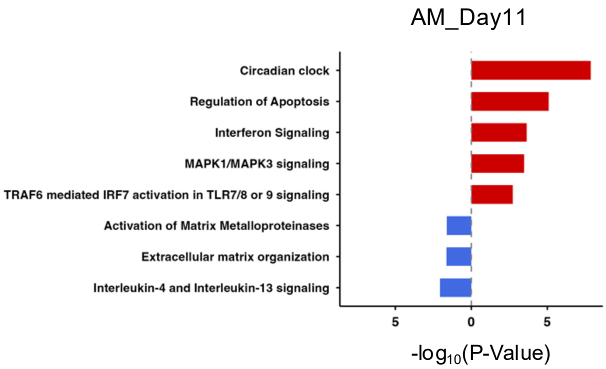
AM_Day11
Circadian clock
Regulation of Apoptosis
Interferon Signaling
MAPK1/MAPK3 signaling
TRAF6 mediated IRF7 activation in TLR7/8 or 9 signaling
Activation of Matrix Metalloproteinases
Extracellular matrix organization
Interleukin-4 and Interleukin-13 signaling
5
0
5
-log10(P-Value)

B

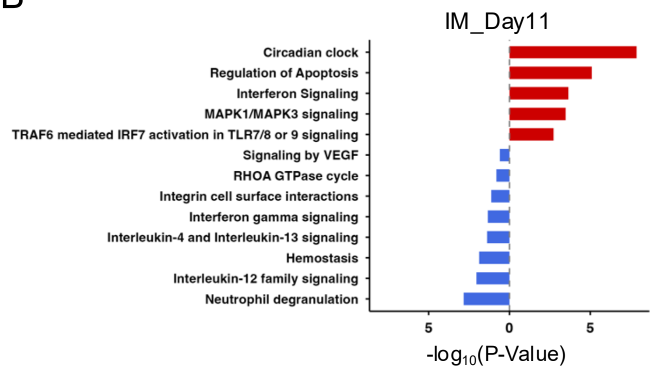
IM_Day11
Circadian clock
Regulation of Apoptosis
Interferon Signaling
MAPK1/MAPK3 signaling
TRAF6 mediated IRF7 activation in TLR7/8 or 9 signaling
Signaling by VEGF
RHOA GTPase cycle
Integrin cell surface interactions
Interferon gamma signaling
Interleukin-4 and Interleukin-13 signaling
Hemostasis
Interleukin-12 family signaling
Neutrophil degranulation
5
0
5
-log10(P-Value)

Sup Fig. 6

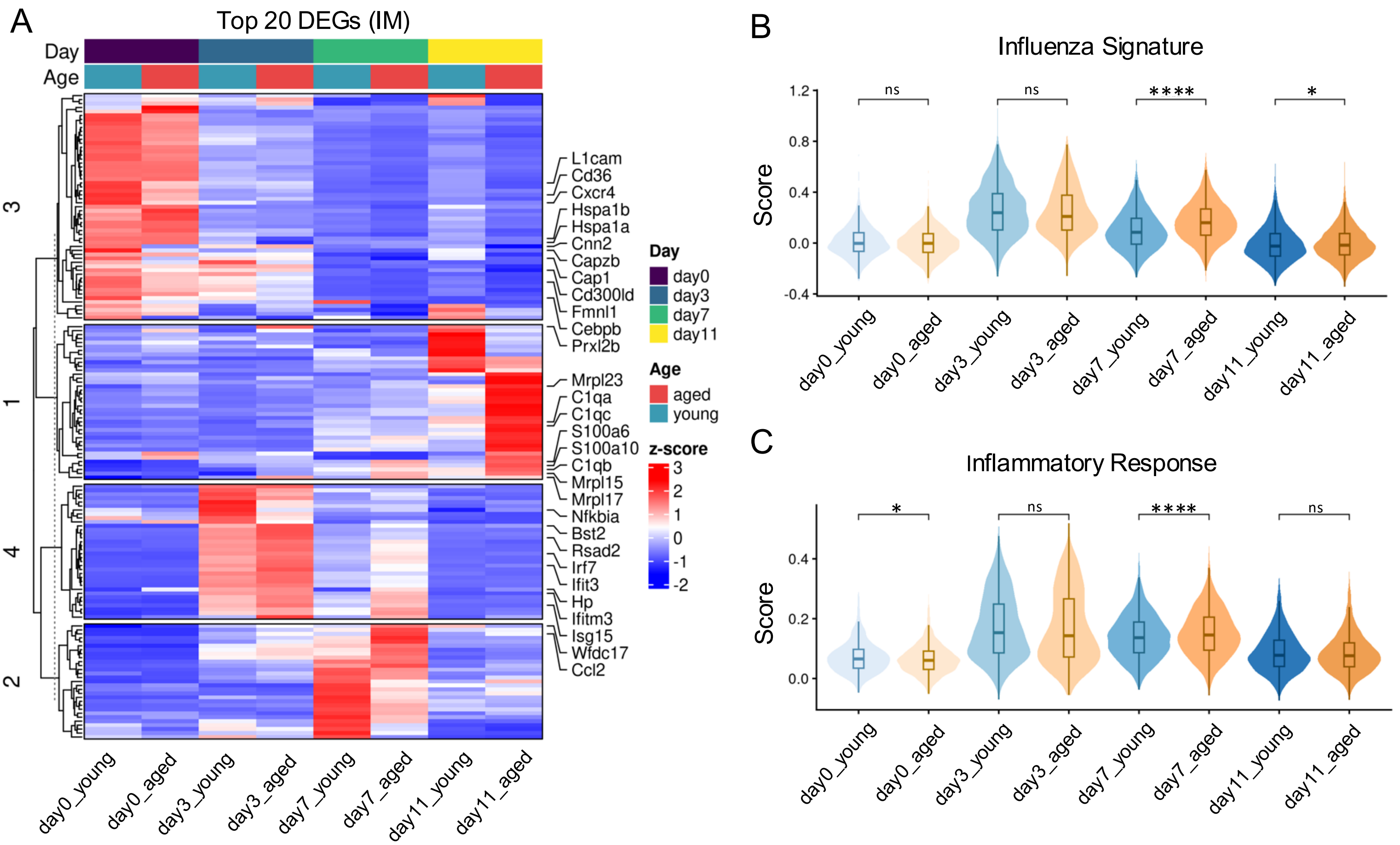

A
Top 20 DEGs (IM)
Day
Age
3
1
4
2
L1cam
Cd36
Cxcr4
Hspa1b
Hspa1a
Cnn2
Capzb
Cap1
Cd300ld
Fmnl1
Cebpb
Prxl2b
Mrpl23
C1qa
C1qc
S100a6
S100a10
C1qb
Mrpl15
Mrpl17
Nfkbia
Bst2
Rsad2
Irf7
Ifit3
Hp
Ifitm3
Isg15
Wfdc17
Ccl2
Day
day0
day3
day7
day11
Age
aged
young
z-score
3
2
1
0
-1
-2
day0_young
day0_aged
day3_young
day3_aged
day7_young
day7_aged
day11_young
day11_aged
B
Influenza Signature
Score
1.2
0.8
0.4
0.0
-0.4
ns
ns
****
*
day0_young
day0_aged
day3_young
day3_aged
day7_young
day7_aged
day11_young
day11_aged
C
Inflammatory Response
Score
0.4
0.2
0.0
*
ns
****
ns
day0_young
day0_aged
day3_young
day3_aged
day7_young
day7_aged
day11_young
day11_aged

Sup Fig. 7

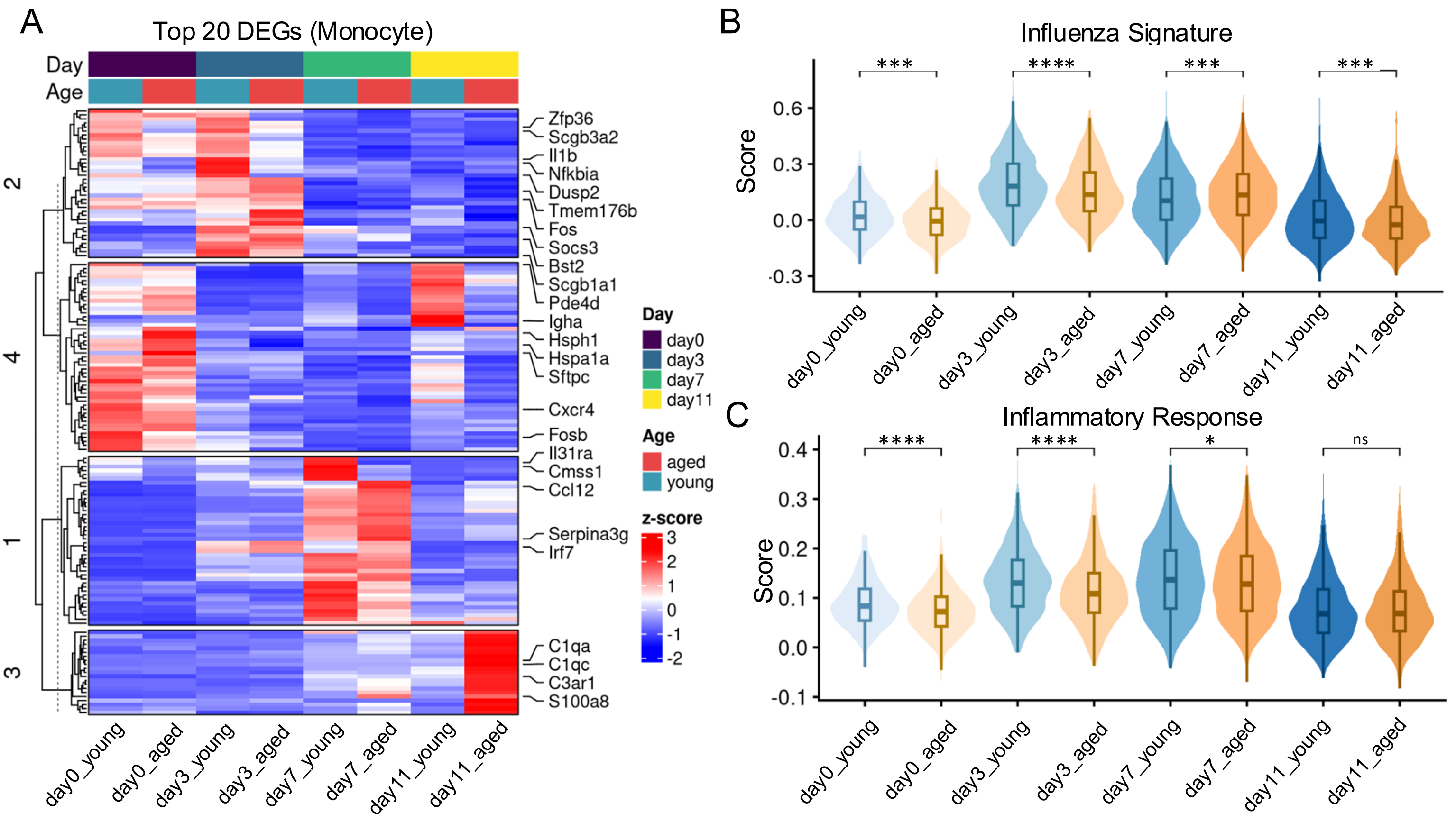

A
Top 20 DEGs (Monocyte)
Day
Age
2
4
1
3
Zfp36
Scgb3a2
Il1b
Nfkbia
Dusp2
Tmem176b
Fos
Socs3
Bst2
Scgb1a1
Pde4d
Igha
Hsph1
Hspa1a
Sftpc
Cxcr4
Fosb
Il31ra
Cmss1
Ccl12
Serpina3g
Irf7
C1qa
C1qc
C3ar1
S100a8
day0_young
day0_aged
day3_young
day3_aged
day7_young
day7_aged
day11_young
day11_aged
Day
day0
day3
day7
day11
Age
aged
young
z-score
3
2
1
0
-1
-2
B
Influenza Signature
Score
0.6
0.3
0.0
-0.3
***
****
***
***
C
Inflammatory Response
Score
0.4
0.3
0.2
0.1
0.0
-0.1
****
****
*
ns

Sup Fig. 8

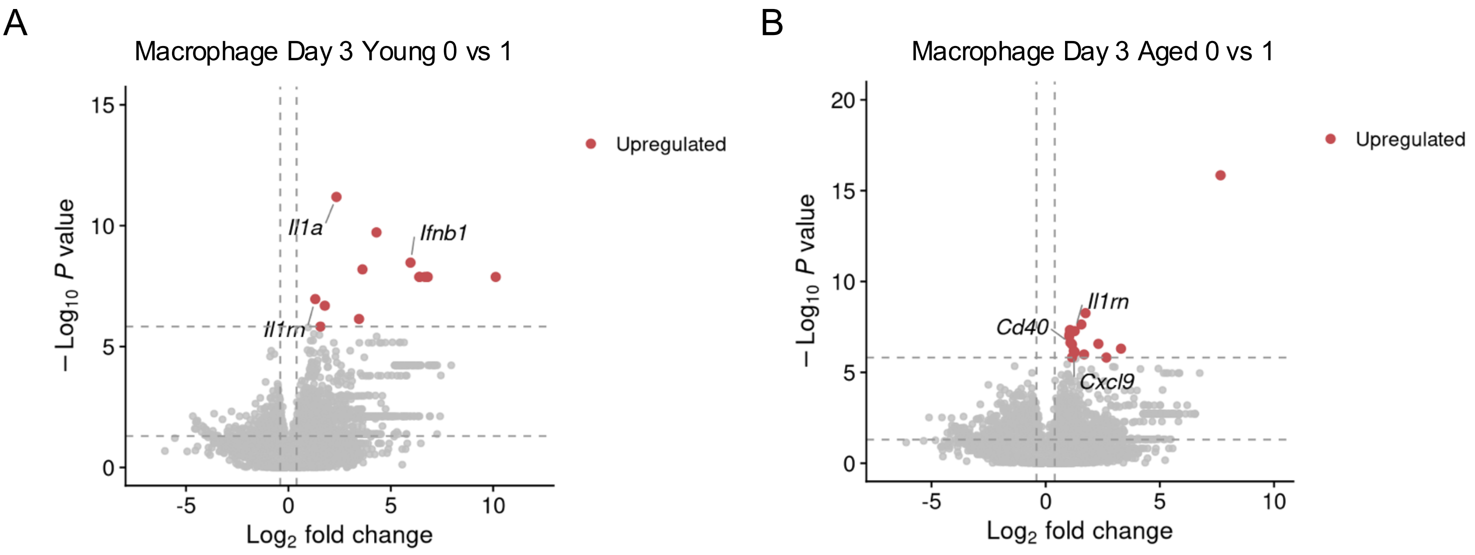
A
Macrophage Day 3 Young 0 vs 1
Upregulated
Il1a
Ifnb1
Il1rn
− Log10 P value
Log2 fold change
B
Macrophage Day 3 Aged 0 vs 1
Upregulated
Il1rn
Cd40
Cxcl9
− Log10 P value
Log2 fold change

Sup Fig. 9

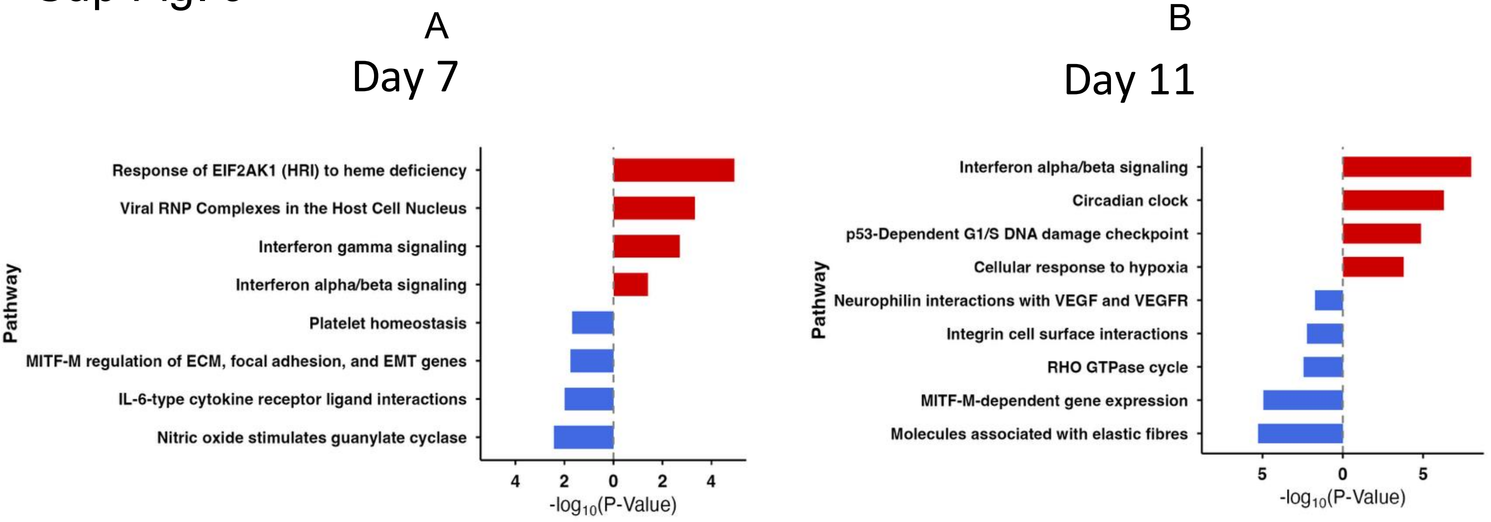
A
Day 7
Response of EIF2AK1 (HRI) to heme deficiency
Viral RNP Complexes in the Host Cell Nucleus
Interferon gamma signaling
Interferon alpha/beta signaling
Platelet homeostasis
MITF-M regulation of ECM, focal adhesion, and EMT genes
IL-6-type cytokine receptor ligand interactions
Nitric oxide stimulates guanylate cyclase
Pathway
4 2 0 2 4
-log10(P-Value)
B
Day 11
Interferon alpha/beta signaling
Circadian clock
p53-Dependent G1/S DNA damage checkpoint
Cellular response to hypoxia
Neurophilin interactions with VEGF and VEGFR
Integrin cell surface interactions
RHO GTPase cycle
MITF-M-dependent gene expression
Molecules associated with elastic fibres
Pathway
5 0 5
-log10(P-Value)

### Sup Fig. 10

A

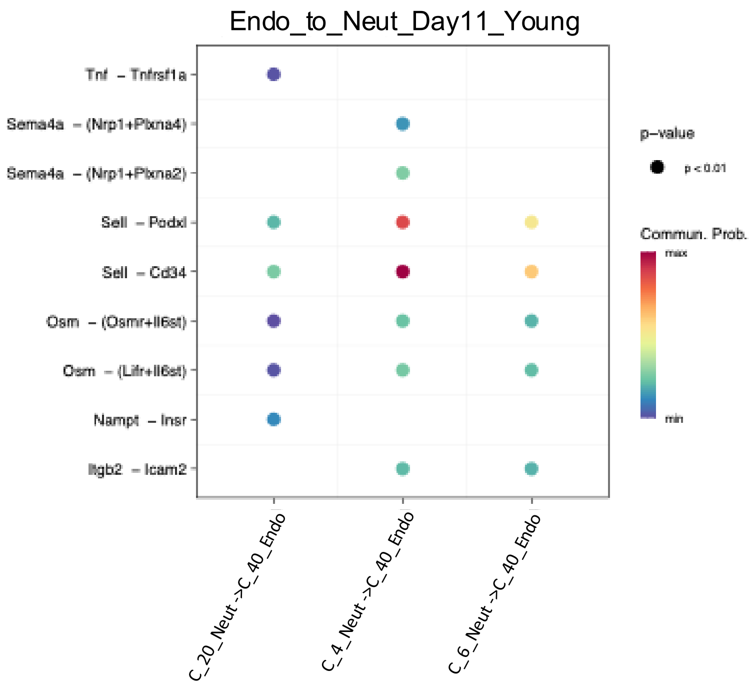
Endo_to_Neut_Day11_Young
Tnf – Tnfrsf1a
Sema4a – (Nrp1+Plxna4)
Sema4a – (Nrp1+Plxna2)
Sell – Podxl
Sell – Cd34
Osm – (Osmr+Il6st)
Osm – (Lifr+Il6st)
Nampt – Insr
Itgb2 – Icam2
C_20_Neut->C_40_Endo
C_4_Neut->C_40_Endo
C_6_Neut->C_40_Endo
p–value
p < 0.01
Commun. Prob.
max
min

B

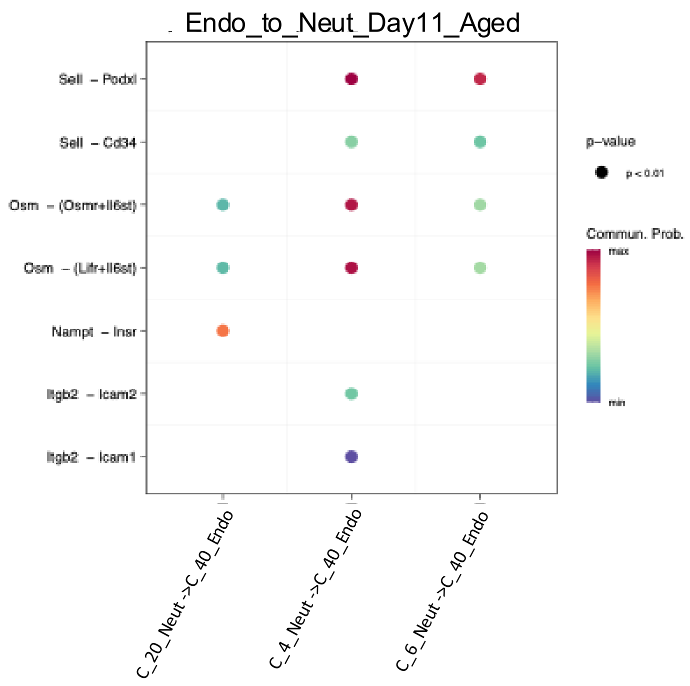
Endo_to_Neut_Day11_Aged
Sell – Podxl
Sell – Cd34
Osm – (Osmr+Il6st)
Osm – (Lifr+Il6st)
Nampt – Insr
Itgb2 – Icam2
Itgb2 – Icam1
C_20_Neut->C_40_Endo
C_4_Neut->C_40_Endo
C_6_Neut->C_40_Endo
p–value
p < 0.01
Commun. Prob.
max
min

Sup Fig. 11

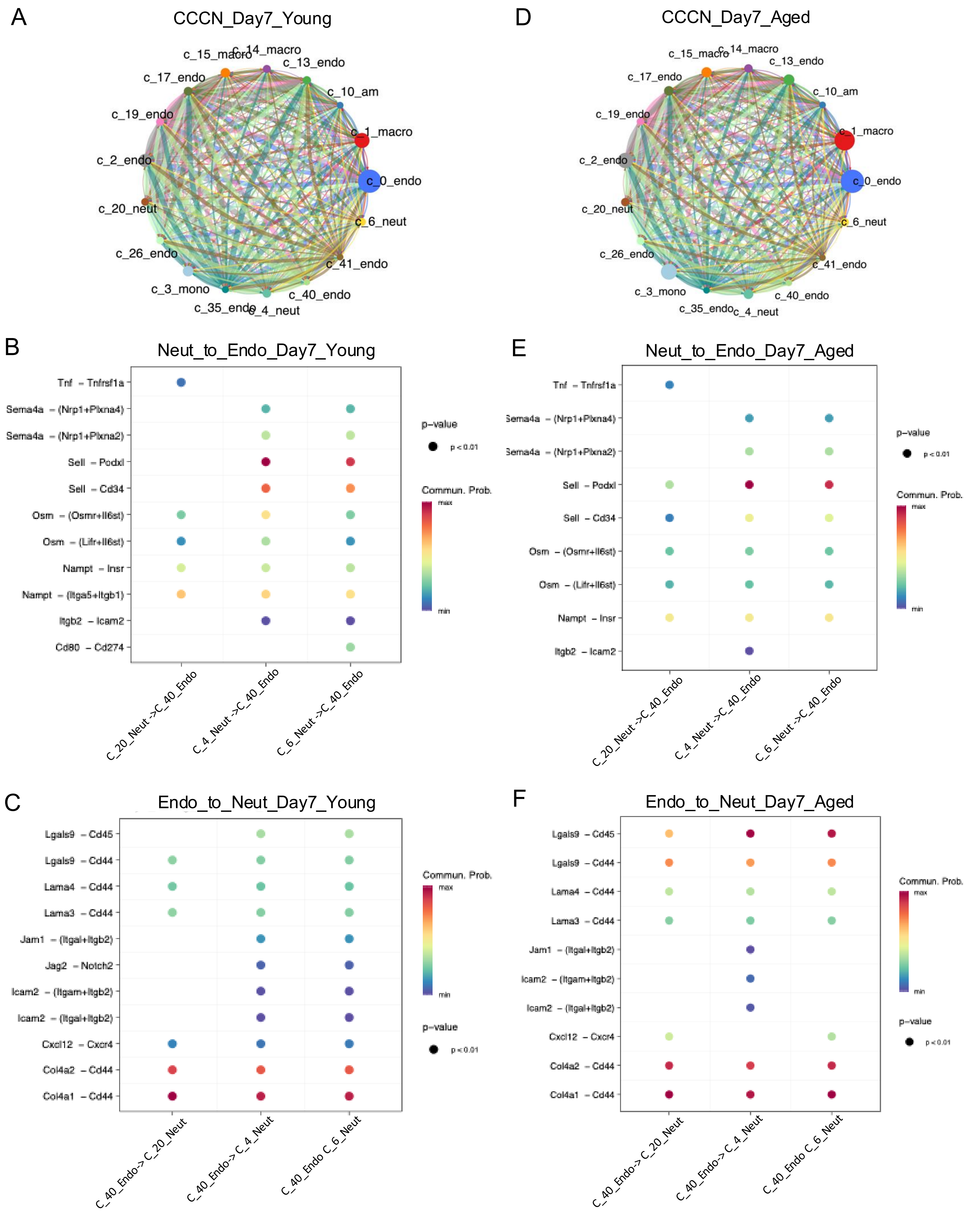

A
CCCN_Day7_Young
c_15_macro
c_14_macro
c_13_endo
c_17_endo
c_10_am
c_19_endo
c_1_macro
c_2_endo
c_0_endo
c_20_neut
c_6_neut
c_26_endo
c_41_endo
c_3_mono
c_40_endo
c_35_endo
c_4_neut
D
CCCN_Day7_Aged
c_15_macro
c_14_macro
c_13_endo
c_17_endo
c_10_am
c_19_endo
c_1_macro
c_2_endo
c_0_endo
c_20_neut
c_6_neut
c_26_endo
c_41_endo
c_3_mono
c_40_endo
c_35_endo
c_4_neut
B
Neut_to_Endo_Day7_Young
Tnf – Tnfrsf1a
Sema4a – (Nrp1+Plxna4)
Sema4a – (Nrp1+Plxna2)
Sell – Podxl
Sell – Cd34
Osm – (Osmr+Il6st)
Osm – (Lifr+Il6st)
Nampt – Insr
Nampt – (Itga5+Itgb1)
Itgb2 – Icam2
Cd80 – Cd274
C_20_Neut->C_40_Endo
C_4_Neut->C_40_Endo
C_6_Neut->C_40_Endo
p-value
p < 0.01
Commun. Prob.
max
min
E
Neut_to_Endo_Day7_Aged
Tnf – Tnfrsf1a
Sema4a – (Nrp1+Plxna4)
Sema4a – (Nrp1+Plxna2)
Sell – Podxl
Sell – Cd34
Osm – (Osmr+Il6st)
Osm – (Lifr+Il6st)
Nampt – Insr
Itgb2 – Icam2
C_20_Neut->C_40_Endo
C_4_Neut->C_40_Endo
C_6_Neut->C_40_Endo
p-value
p < 0.01
Commun. Prob.
max
min
C
Endo_to_Neut_Day7_Young
Lgals9 – Cd45
Lgals9 – Cd44
Lama4 – Cd44
Lama3 – Cd44
Jam1 – (Itgal+Itgb2)
Jag2 – Notch2
Icam2 – (Itgam+Itgb2)
Icam2 – (Itgal+Itgb2)
Cxcl12 – Cxcr4
Col4a2 – Cd44
Col4a1 – Cd44
C_40_Endo->C_20_Neut
C_40_Endo->C_4_Neut
C_40_Endo C_6_Neut
Commun. Prob.
max
min
p-value
p < 0.01
F
Endo_to_Neut_Day7_Aged
Lgals9 – Cd45
Lgals9 – Cd44
Lama4 – Cd44
Lama3 – Cd44
Jam1 – (Itgal+Itgb2)
Icam2 – (Itgam+Itgb2)
Icam2 – (Itgal+Itgb2)
Cxcl12 – Cxcr4
Col4a2 – Cd44
Col4a1 – Cd44
C_40_Endo->C_20_Neut
C_40_Endo->C_4_Neut
C_40_Endo C_6_Neut
Commun. Prob.
max
min
p-value
p < 0.01

### Sup Fig. 12

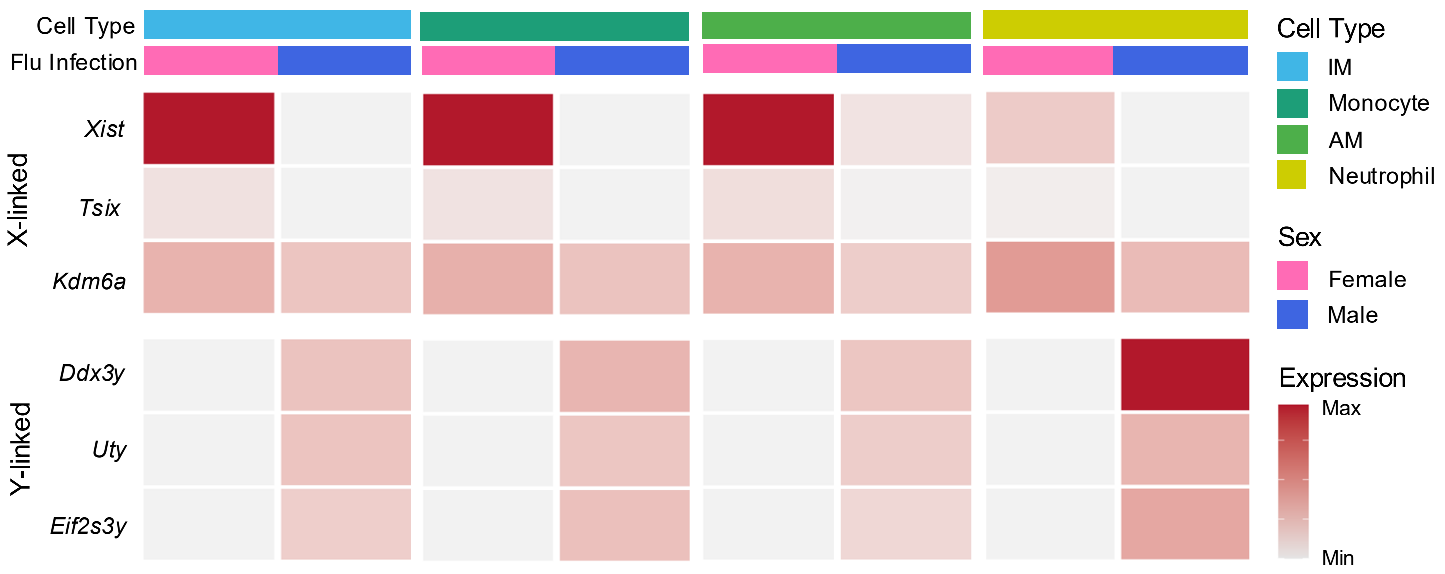

Cell Type
Flu Infection
X-linked
Xist
Tsix
Kdm6a
Y-linked
Ddx3y
Uty
Eif2s3y
Cell Type
IM
Monocyte
AM
Neutrophil
Sex
Female
Male
Expression
Max
Min

Sup Fig. 13

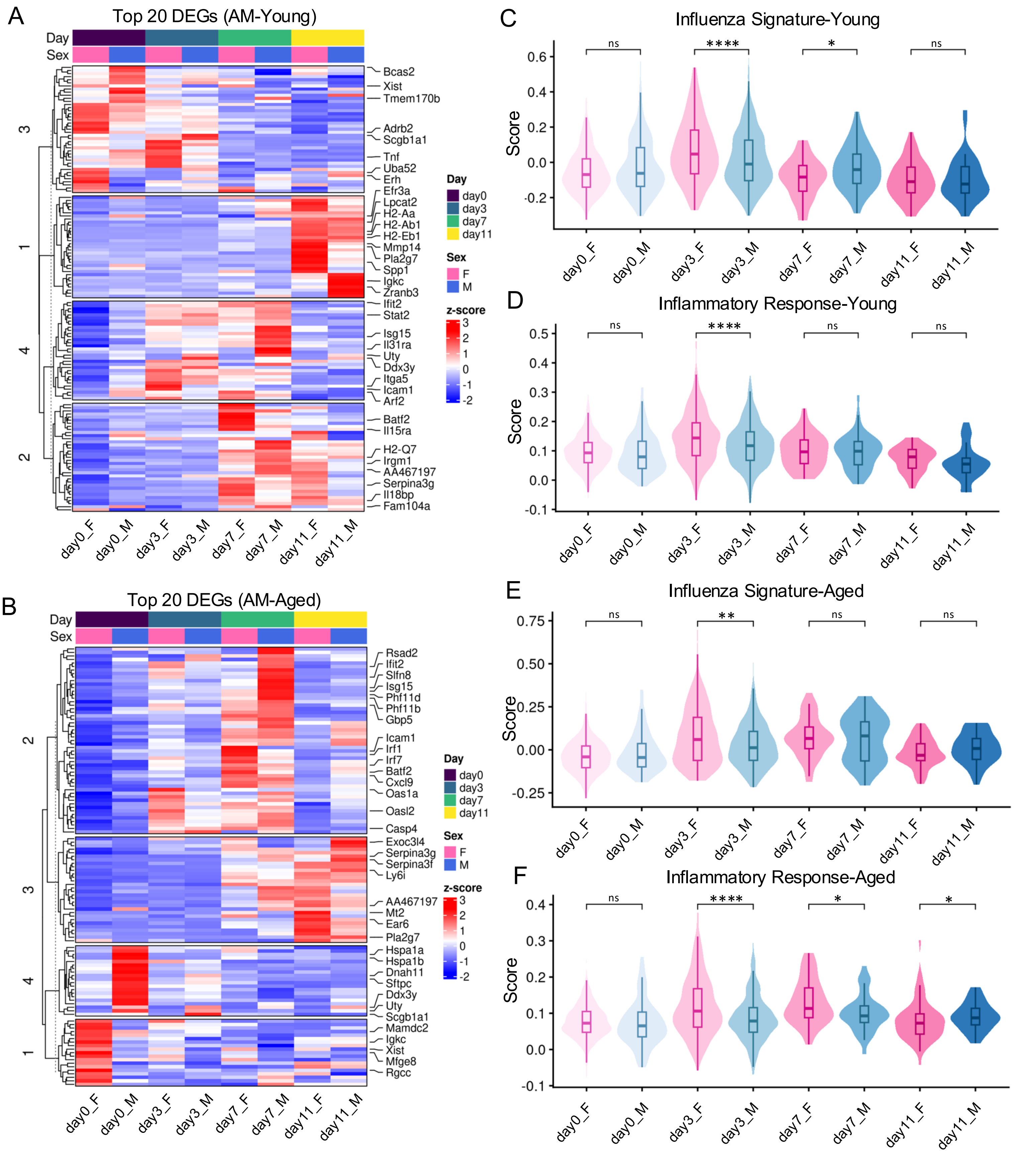

A
Top 20 DEGs (AM-Young)
Day
Sex
3
1
4
2
Bcas2
Xist
Tmem170b
Adrb2
Scgb1a1
Tnf
Uba52
Erh
Efr3a
Lpcat2
H2-Aa
H2-Ab1
H2-Eb1
Mmp14
Pla2g7
Spp1
Igkc
Zranb3
Ifit2
Stat2
Isg15
Il31ra
Uty
Ddx3y
Itga5
Icam1
Arf2
Batf2
Il15ra
H2-Q7
Irgm1
AA467197
Serpina3g
Il18bp
Fam104a
Day
day0
day3
day7
day11
Sex
F
M
z-score
3
2
1
0
-1
-2
day0_F
day0_M
day3_F
day3_M
day7_F
day7_M
day11_F
day11_M
B
Top 20 DEGs (AM-Aged)
Day
Sex
2
3
4
1
Rsad2
Ifit2
Slfn8
Isg15
Phf11d
Phf11b
Gbp5
Icam1
Irf1
Irf7
Batf2
Cxcl9
Oas1a
Oasl2
Casp4
Exoc3l4
Serpina3g
Serpina3f
Ly6i
AA467197
Mt2
Ear6
Pla2g7
Hspa1a
Hspa1b
Dnah11
Sftpc
Ddx3y
Uty
Scgb1a1
Mamdc2
Igkc
Xist
Mfge8
Rgcc
Day
day0
day3
day7
day11
Sex
F
M
z-score
3
2
1
0
-1
-2
day0_F
day0_M
day3_F
day3_M
day7_F
day7_M
day11_F
day11_M
C
Influenza Signature-Young
ns
****
*
ns
Score
0.6
0.4
0.2
0.0
-0.2
day0_F
day0_M
day3_F
day3_M
day7_F
day7_M
day11_F
day11_M
D
Inflammatory Response-Young
ns
****
ns
ns
Score
0.5
0.4
0.3
0.2
0.1
0.0
-0.1
day0_F
day0_M
day3_F
day3_M
day7_F
day7_M
day11_F
day11_M
E
Influenza Signature-Aged
ns
**
ns
ns
Score
0.75
0.50
0.25
0.00
-0.25
day0_F
day0_M
day3_F
day3_M
day7_F
day7_M
day11_F
day11_M
F
Inflammatory Response-Aged
ns
****
*
*
Score
0.4
0.3
0.2
0.1
0.0
-0.1
day0_F
day0_M
day3_F
day3_M
day7_F
day7_M
day11_F
day11_M

Sup Fig. 14

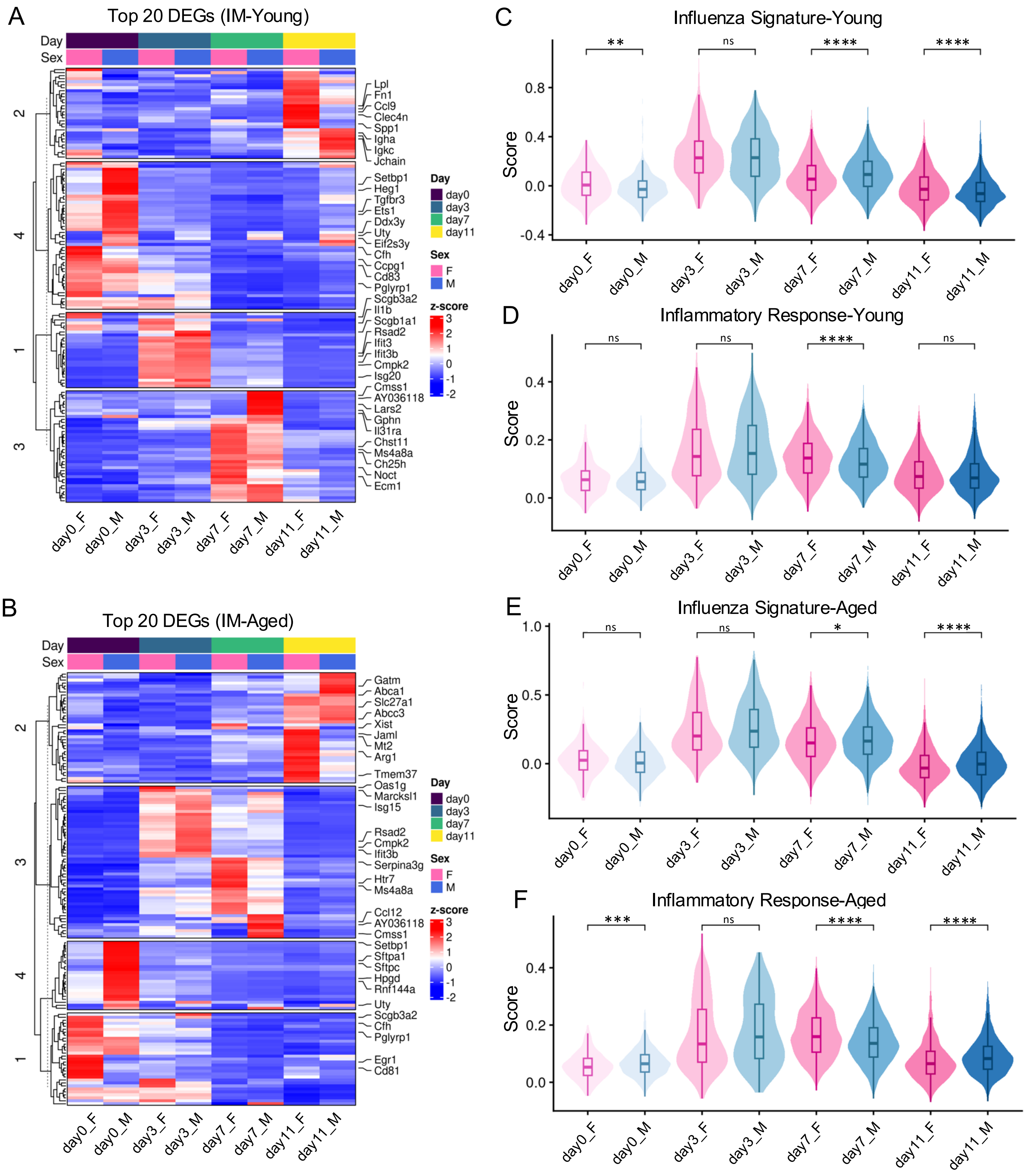
A
Top 20 DEGs (IM-Young)
Day
Sex
Lpl
Fn1
Ccl9
Clec4n
Spp1
Igha
Igkc
Jchain
Setbp1
Heg1
Tgfbr3
Ets1
Ddx3y
Uty
Eif2s3y
Cfh
Ccpg1
Cd83
Pglyrp1
Scgb3a2
Il1b
Scgb1a1
Rsad2
Ifit3
Ifit3b
Cmpk2
Isg20
Cmss1
AY036118
Lars2
Gphn
Il31ra
Chst11
Ms4a8a
Ch25h
Noct
Ecm1
2
4
1
3
day0_F
day0_M
day3_F
day3_M
day7_F
day7_M
day11_F
day11_M
Day
day0
day3
day7
day11
Sex
F
M
z-score
3
2
1
0
-1
-2
B
Top 20 DEGs (IM-Aged)
Gatm
Abca1
Slc27a1
Abcc3
Xist
Jaml
Mt2
Arg1
Tmem37
Oas1g
Marcksl1
Isg15
Rsad2
Cmpk2
Ifit3b
Serpina3g
Htr7
Ms4a8a
Ccl12
AY036118
Cmss1
Setbp1
Sftpa1
Sftpc
Hpgd
Rnf144a
Uty
Scgb3a2
Cfh
Pglyrp1
Egr1
Cd81
C
Influenza Signature-Young
Score
**
ns
****
****
D
Inflammatory Response-Young
ns
ns
****
ns
E
Influenza Signature-Aged
ns
ns
*
****
F
Inflammatory Response-Aged
***
ns
****
****

Sup Fig. 15

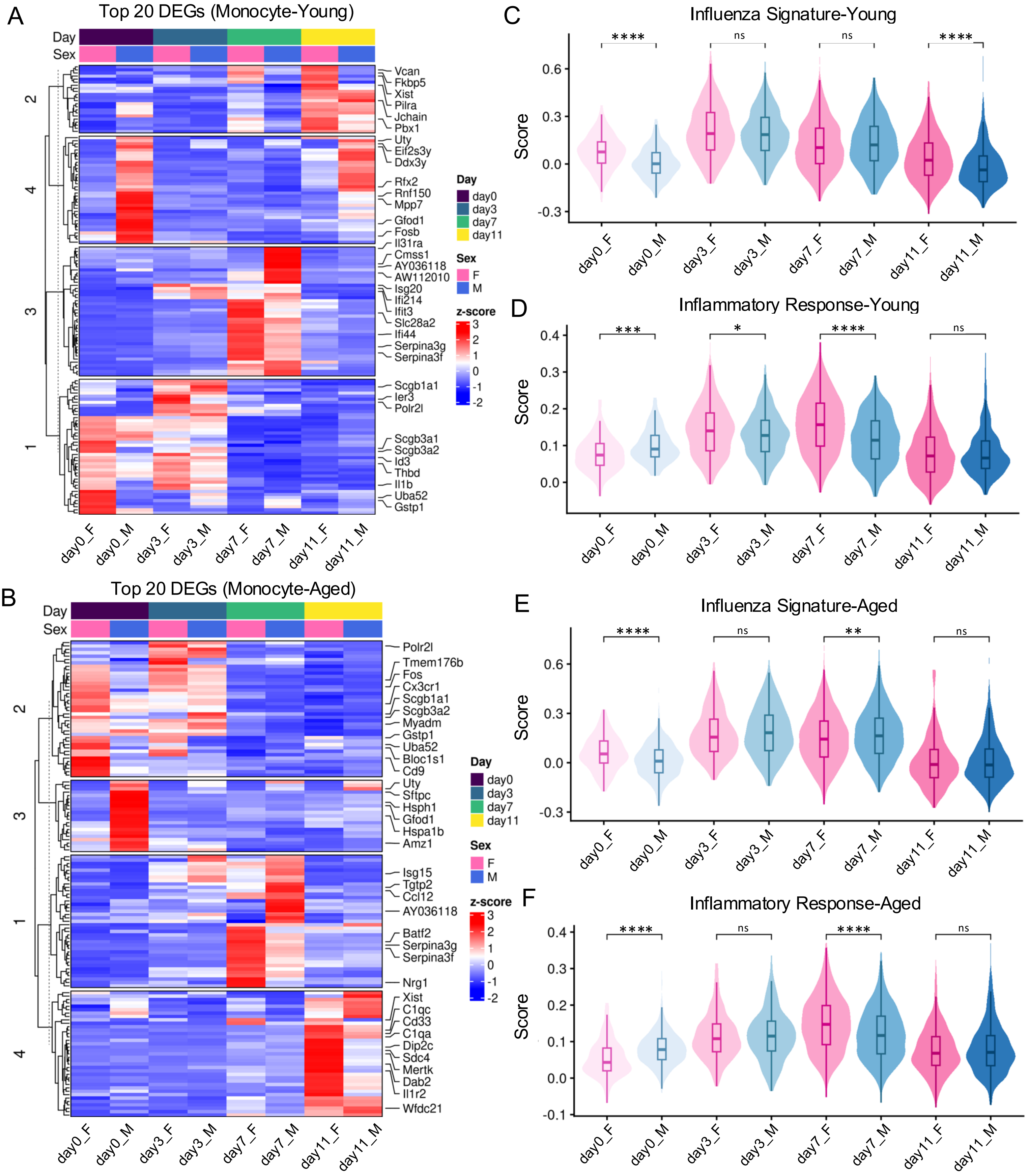
A
Top 20 DEGs (Monocyte-Young)
Day
Sex
2
4
3
1
Vcan
Fkbp5
Xist
Pilra
Jchain
Pbx1
Uty
Eif2s3y
Ddx3y
Rfx2
Rnf150
Mpp7
Gfod1
Fosb
Il31ra
Cmss1
AY036118
AW112010
Isg20
Ifi214
Ifit3
Slc28a2
Ifi44
Serpina3g
Serpina3f
Scgb1a1
Ier3
Polr2l
Scgb3a1
Scgb3a2
Id3
Thbd
Il1b
Uba52
Gstp1
day0_F
day0_M
day3_F
day3_M
day7_F
day7_M
day11_F
day11_M
Day
day0
day3
day7
day11
Sex
F
M
z-score
3
2
1
0
-1
-2
B
Top 20 DEGs (Monocyte-Aged)
Day
Sex
2
3
1
4
Polr2l
Tmem176b
Fos
Cx3cr1
Scgb1a1
Scgb3a2
Myadm
Gstp1
Uba52
Bloc1s1
Cd9
Uty
Sftpc
Hsph1
Gfod1
Hspa1b
Amz1
Isg15
Tgtp2
Ccl12
AY036118
Batf2
Serpina3g
Serpina3f
Nrg1
Xist
C1qc
Cd33
C1qa
Dip2c
Sdc4
Mertk
Dab2
Il1r2
Wfdc21
day0_F
day0_M
day3_F
day3_M
day7_F
day7_M
day11_F
day11_M
Day
day0
day3
day7
day11
Sex
F
M
z-score
3
2
1
0
-1
-2
C
Influenza Signature-Young
Score
****
ns
ns
****
0.6
0.3
0.0
-0.3
day0_F
day0_M
day3_F
day3_M
day7_F
day7_M
day11_F
day11_M
D
Inflammatory Response-Young
Score
***
*
****
ns
0.4
0.3
0.2
0.1
0.0
day0_F
day0_M
day3_F
day3_M
day7_F
day7_M
day11_F
day11_M
E
Influenza Signature-Aged
Score
****
ns
**
ns
0.6
0.3
0.0
-0.3
day0_F
day0_M
day3_F
day3_M
day7_F
day7_M
day11_F
day11_M
F
Inflammatory Response-Aged
Score
****
ns
****
ns
0.4
0.3
0.2
0.1
0.0
-0.1
day0_F
day0_M
day3_F
day3_M
day7_F
day7_M
day11_F
day11_M

Sup Fig. 16

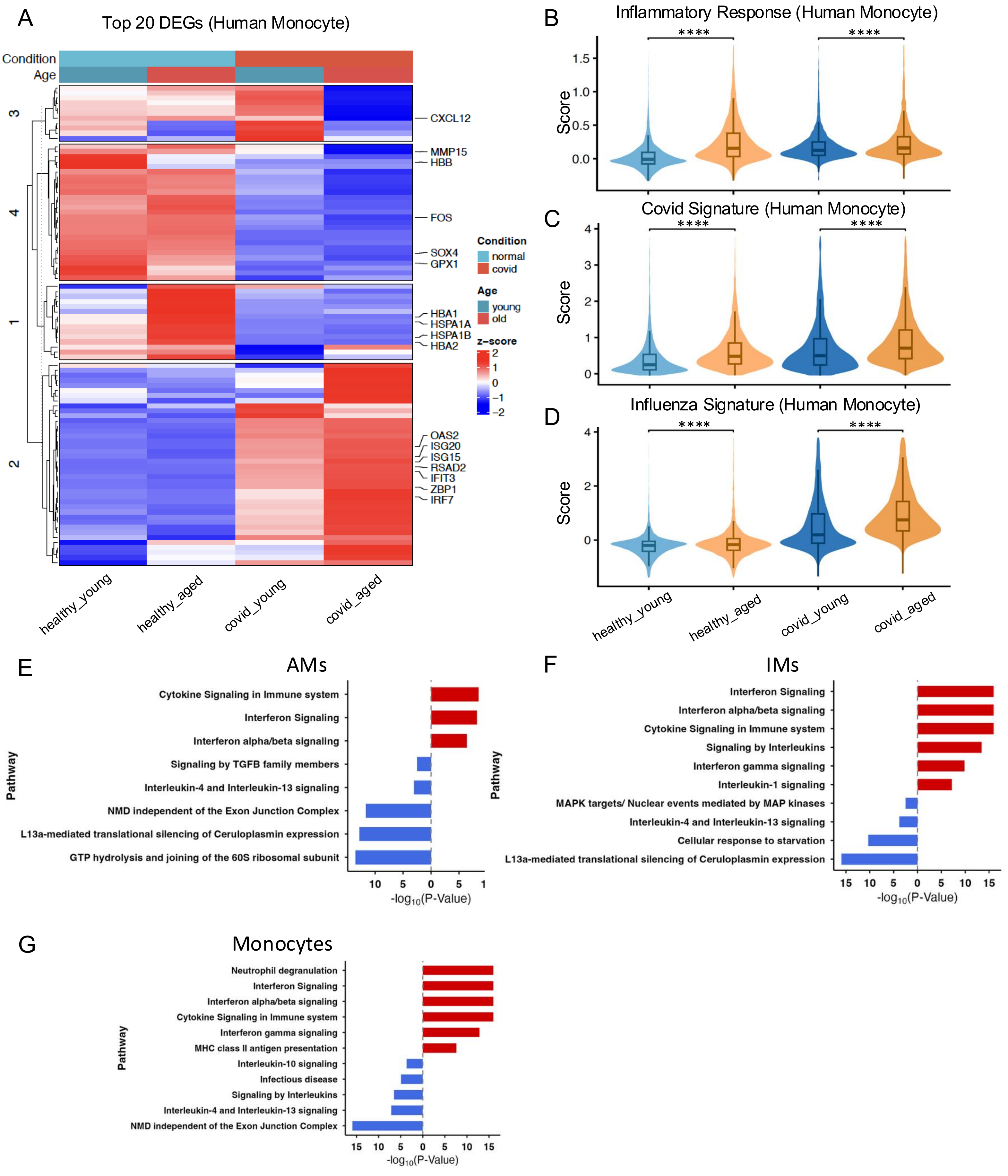
A
Top 20 DEGs (Human Monocyte)
Condition
Age
3
4
1
2
CXCL12
MMP15
HBB
FOS
SOX4
GPX1
HBA1
HSPA1A
HSPA1B
HBA2
OAS2
ISG20
ISG15
RSAD2
IFIT3
ZBP1
IRF7
Condition
normal
covid
Age
young
old
z-score
2
1
0
−1
−2
healthy_young
healthy_aged
covid_young
covid_aged
B
Inflammatory Response (Human Monocyte)
****
****
Score
1.5
1.0
0.5
0.0
C
Covid Signature (Human Monocyte)
****
****
Score
4
3
2
1
0
D
Influenza Signature (Human Monocyte)
****
****
Score
4
2
0
healthy_young
healthy_aged
covid_young
covid_aged
E
AMs
Pathway
Cytokine Signaling in Immune system
Interferon Signaling
Interferon alpha/beta signaling
Signaling by TGFB family members
Interleukin-4 and Interleukin-13 signaling
NMD independent of the Exon Junction Complex
L13a-mediated translational silencing of Ceruloplasmin expression
GTP hydrolysis and joining of the 60S ribosomal subunit
10 5 0 5 1
-log10(P-Value)
F
IMs
Pathway
Interferon Signaling
Interferon alpha/beta signaling
Cytokine Signaling in Immune system
Signaling by Interleukins
Interferon gamma signaling
Interleukin-1 signaling
MAPK targets/ Nuclear events mediated by MAP kinases
Interleukin-4 and Interleukin-13 signaling
Cellular response to starvation
L13a-mediated translational silencing of Ceruloplasmin expression
15 10 5 0 5 10 15
-log10(P-Value)
G
Monocytes
Pathway
Neutrophil degranulation
Interferon Signaling
Interferon alpha/beta signaling
Cytokine Signaling in Immune system
Interferon gamma signaling
MHC class II antigen presentation
Interleukin-10 signaling
Infectious disease
Signaling by Interleukins
Interleukin-4 and Interleukin-13 signaling
NMD independent of the Exon Junction Complex
15 10 5 0 5 10 15
-log10(P-Value)
